# Environment-organism feedbacks drive changes in ecological interactions

**DOI:** 10.1101/2023.10.31.565024

**Authors:** Oliver J. Meacock, Sara Mitri

## Abstract

Ecological interactions, the impact of one organism on the growth and death of another, underpin our understanding of the long-term composition and the functional properties of communities. In recent years, the context-dependency of interactions – their tendency to change values in different environments, locations and at different times – has become an increasingly important theme in ecological research. However, an overarching theoretical assumption has been that external environmental factors are responsible for driving these changes. Here, we derive a theoretical interaction framework which teases apart the separate roles played by these extrinsic environmental inputs and the intrinsic environmental changes driven by organisms within the environment itself. At the heart of our theory is the ‘instantaneous interaction’, a quantity that captures the feedback between environmental composition and the growth of organisms within it. In the limit that intrinsic, organism-driven environmental change dominates over external drivers, we find that this feedback can give rise to temporal and spatial context-dependencies as organisms modify the environment over time and/or space. We use small synthetic microbial communities as model ecosystems to demonstrate the power of this framework, using it to predict time-dependent intra-specific interactions in a toxin degradation system and to relate time and spatial dependencies in crossfeeding communities. Our framework helps to explain the ubiquity of interaction context-dependencies in systems where population changes are driven by environmental changes – such as microbial communities – by placing the environment on an equal theoretical footing as the organisms that inhabit it.

## 1 Introduction

Interactions between organisms – the impact of one species on the fitness or growth rate of another^1,2^ – are one of the most consistent themes in ecology. Originally, theoretical ecologists conceived of interactions as fixed quantities,^3–7^ allowing them to be assembled into frameworks that predict community-level properties based on elementary, pairwise interaction estimates. ^5,8^ This assumption has, however, proven to be shaky in recent decades; interactions are now known to vary depending on the environment in which they are measured^9–13^ and the time at which they are measured, ^13–15^ and spatial structure in multi-species communities is at least qualitatively understood to influence inter-species interactions.^16^ Such context-dependencies substantially complicate bottom-up attempts to predict community-level outcomes based on assemblages of elementary interaction measurements, as the context in which the interaction between a pair of organisms is measured may be very different from that in which the assembled community resides. Recent results from microbial ecology support this view, showing that pairwise measurements often fail to capture community-level outcomes.^17,18^ To resolve these issues, we must first understand how context-dependencies arise and, if possible, predict them.

Numerous underlying mechanisms of context-dependency are known. Over evolutionary timescales, interaction values can change in response to environmentally-dependent selective pressures that modify the strategies of organisms.^19^ Yet even over timescales where strategies remain fixed, environmental factors can also influence interactions. While external, so-called *allogenic* factors such as climactic change are one such component of environmental change,^20,21^ organisms can themselves influence their local environment, especially in sessile communities. Such *autogenic* environmental changes can mediate interactions when they modify the growth rate of surrounding organisms, effectively setting up a feedback loop between community composition and environmental composition. For example, reduction of local salinity by nurse plants in salt marshes can allow establishment of salt-sensitive species,^22^ while the detoxification of environmental compounds by partners allows toxin-sensitive species to grow to higher densities in microbial culture. ^9^ In these cases, net positive interactions result as long as autogenic mechanisms that increase growth rates (such as stress buffering) outweigh mechanisms that decrease growth (such as nutrient competition). This balance shifts between different underlying environments, resulting in a switch from net negative to net positive interactions along stress gradients – an example of an environmental context dependency. ^11,23–25^ Despite this qualitative understanding of the importance of autogenic processes in driving interaction changes however, we lack corresponding theory to predict these changes and generalise the phenomena to other systems.

In this manuscript, we provide a theoretical basis for predicting environmental, temporal and spatial context dependencies based on the feedback between the growth rates of organisms in different environments and their autogenic impact on their environment. Our approach builds on classical Consumer-Resource (CR) models, which explicitly represent the mechanisms of resource uptake through which many organisms compete; ^7,26^ we adopt a generalised version of this approach that incorporates inhibitory environmental factors and positive impacts of organisms on their environment (*e.g.* secretion of compounds), which we refer to as the Environment-Organism (EO) framework. ^27^ We show that a quantity we call the ‘instantaneous interaction’ – the environmentally-dependent effect that one species has on the growth of another – arises naturally out of the fundamental equations describing EO systems. Communities in environments subject to purely autogenic change sweep out trajectories in the space of possible environments over time or space, and this changing environmental context leads to predictable variations in the instantaneous interaction. In the most dramatic cases, this can lead to switching of the sign of interactions, from positive to negative or vice versa. We then experimentally verify these predictions using small microbial communities for which allogenic mechanisms of environmental change can be eliminated, allowing us to isolate the role of the autogenic processes. Our work thus provides a basis for disentangling autogenic and allogenic contributors to context-dependencies.

## 2 Results

### 2.1 A theoretical Environment-Organism interaction framework explains multiple context-dependencies

We begin by considering general EO systems for which complex interactions can be decomposed into elementary components, each mediated by a single environmental factor. Our goal will be to write a general expression for population dynamics containing an interaction term that directly links the environmental context of the system to the measured interactions between species.

EO systems can be modelled by breaking them into three parts: ^26,28,29^ firstly the *impact function* of a species *β*, ***f****_β_*(***r***) describes the rate at which one unit of *β* modifies its environment *i.e.* the autogenic component of environmental change. We denote this environment with the vector ***r***, which for the purposes of this manuscript we will mostly take to represent the concentrations of different chemical intermediates (*e.g.* element 1 represents the concentration of glucose, while element 2 represents the concentration of acetate), but may more generally represent quantities such as temperature and light availability. ***r*** defines a position in the ‘environment space’, the set of different possible environmental states. The impact function is itself dependent upon ***r***, allowing it to capture, for example, concentration-dependent uptake of a resource. ***r*** will also be impacted by the second component ***σ***, which represents allogenic processes such as flows of intermediates into or out of the system. We can then write the rate of change of the environment as:

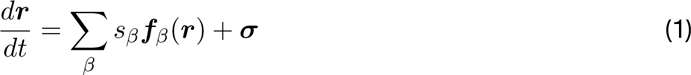

where *s_β_* is the instantaneous abundance of species *β*.

Thirdly, the *sensitivity function g_α_* describes the per-capita growth rate of a species *α* in a particular environment:

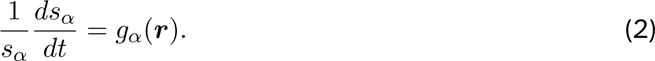

As defined here, these functions are very general, allowing expression of various categories of EO relationship. These include ‘switching’ phenotypes such as diauxy, whereby organisms selectively impact the environmental factor that maximises their growth rate, as well as combinations of essential resources that must be impacted simultaneously during growth. ^26^

The dependence of *g_α_* on ***r*** means that changes to the environment caused by both *α* itself (*β* = *α*, intra-specific interactions) and the other species (*β ̸*= *α*, inter-specific interactions) (Eq. 1) will regulate *α*’s growth rate. Breaking this regulation into the effect mediated by each environmental factor *r_ρ_* individually, we can define ‘elementary’ interactions. These can be categorised into four classes by the combinations of the signs of the impact and sensitivity functions; following recently-defined terminology for metabolic interactions,^29,30^ we refer to these here as enrichment (*β* produces a nutrient that enhances the growth of *α*), depletion (*β* reduces the concentration of a nutrient, impeding the growth of *α*), pollution (*β* produces a toxin that impedes the growth of *α*) and detoxification (*β* decreases the concentration of a toxin of *α* and enhances its growth) (Fig. 1A).

**Figure 1:**
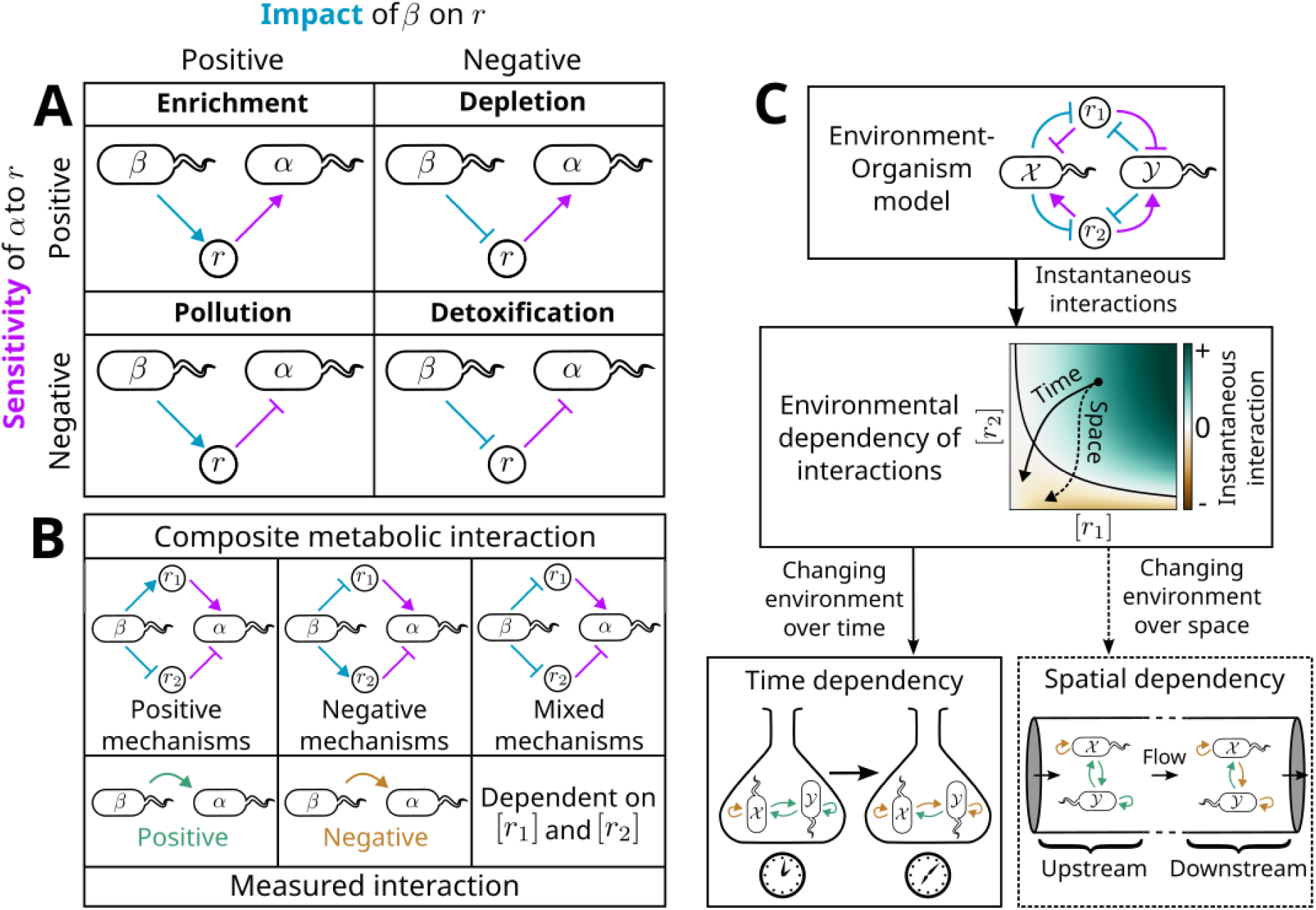
Multiple interaction context-dependencies can be explained with a single theoretical framework. **A** Indirect interactions between organisms break into elementary components when the role of each environmental factor *r_ρ_*is considered separately. ‘Sensitivity functions’ (purple) denote the effect of increasing a factor *r* on the growth rate of a target species *α* (*i.e.* whether it decreases – bar – or increases – arrow – *α*’s growth), while ‘impact functions’ (blue) denote the effect of an effector species *β* on *r* (*i.e.* whether it is increased – arrow – or decreased – bar). Combinations of these functions imply four elementary interaction types: enrichment and detoxification which enhance the growth of *α*, and depletion and pollution which reduce *α*’s growth. ^29,30^ **B** Combinations of elementary interactions yield composite interactions, with the measured interaction’s sign depending on the individual effects of the composed elements. When positive and negative elementary mechanisms are mixed, the measured interaction depends on the balance between the factors – *i.e.* the environmental context. **C** Our framework shows how Environment-Organism (EO) models give rise to an instantaneous interaction that depends on the environment. As the environment changes over time (*e.g.* in batch culture) or over space (*e.g.* in microfluidic channels at steady-state), this environmental dependency in turn gives rise to time and spatial dependencies.

Species can interact via any combination of these elementary interactions, resulting in ‘composite’ metabolic interactions. In general, if all elementary interactions forming a composite interaction mediate a positive growth-rate effect on *β* (enrichment or detoxification) the measured interaction will also be positive, if all mediate a negative growth-rate effect (depletion or pollution) the measured interaction with be negative, and if they have a mixture of positive and negative impacts the sign of the measured interaction will depend on the relative balance of the positive and negative mechanisms, in turn dependent upon the environmental context (Fig. 1B).

We can capture this environmental dependency naturally within the impact/sensitivity function framework. In systems dominated by autogenic mechanisms of environmental change – such that ***σ*** = **0** – it can be shown that (Supplementary Text 1):

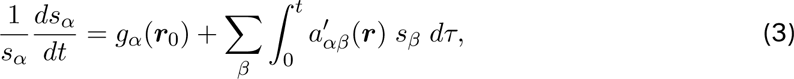

where ***r***_0_ is the initial environmental composition and the integral is taken over the entire history of the system up to the current time *t* (parameterised by *τ*). We will refer to this expression as the closed Environment-Organism (cEO) equation, as the environment is closed with respect to allogenic influences.

A key component of this expression is the *instantaneous interaction* 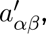 defined as

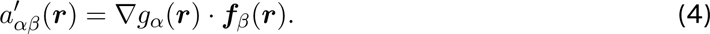

*∇g_α_*(***r***) is the gradient of the sensitivity function, a vector field which denotes the direction in the environment space along which the growth rate of *α* increases most rapidly. The scalar product of this with ***f****_β_*(***r***) (also a vector field) can therefore be naturally interpreted as indicating whether *β* is pulling the environment in a direction that increases (positive 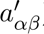) or decreases (negative 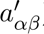) the growth of *α* at a given position in the environment space ***r***. Put simply, this term captures the environmental dependency of the interaction in a given environment. In Supplementary Text 1, we show that the same term arises in systems where autogenic and allogenic mechanisms are in equilibrium, such as chemostats. In such settings, 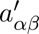 plays an equivalent role to the interaction matrix. ^5^ We can exploit this link to approximate interactions in mixed autogenic/allogenic settings by isolating the equilibrium environment from allogenic influences and measuring the resulting growth rate dynamics of pairs of organisms (Supplementary Text 2).

What do the other parts of this expression tell us about the dynamics of purely autogenic ecosystems? To answer this, it is instructive to compare the cEO equation with the generalised Lotka-Volterra (gLV) equation:

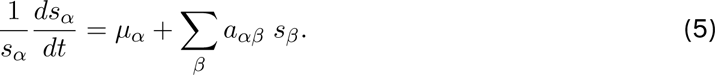

Here, *µ_α_* is *α*’s intrinsic growth rate (*i.e.* its growth in the absence of other species and at low population sizes) and *a_αβ_* is the interaction between *β* and *α*, defined as the population-dependent impact of *β* on the growth rate of *α*. We note some similarities between the two equations: both are expressed in terms of a basal growth rate added to a sum of interaction terms from all species *β* interacting with *α*. However, there are two important distinctions between the notion of interactions in the cEO and gLV frameworks. Firstly, as previously noted, 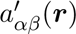 is dependent on the environmental context. Moreover, this environment-dependence is not static – in general, environments under autogenic control trace out some trajectory ***r***(*t*) in the environment space, over which 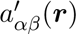 can vary dramatically. Indeed, we will soon see that in many systems it can change sign over time and space. Secondly, interactions in the EO framework are cumulative, arising from the integration of the interaction term over the entire history of the system. This reflects the fact that interactions are mediated via ongoing changes to environmental factors, which take time to be impacted by organisms. We refer to the resulting net impact of *β* on *α*’s growth rate at a given time 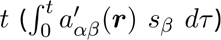 as the *cumulative interaction*.

In the remainder of this manuscript, we illustrate how the environmental dependency of the instantaneous interaction results in temporal interaction dependencies and spatial structure when the environmental context changes autogenically over time and space (Fig. 1C). We further show how these context dependencies can be predicted if underlying interaction mechanisms (represented by appropriate choices of the impact and sensitivity functions) are already known.

### 2.2 Composite interactions can result in time-dependencies of interaction measurements

One of the simplest microbial interactions with mixed positive and negative components is that of a single species *A* interacting negatively with itself via nutrient depletion and positively via detoxification (Fig. 2A). We can simulate such a system using the impact (Fig. 2B) and sensitivity functions (Fig. 2C) for this system from a previously described EO framework ^9^ in which nutrient uptake and toxin impact are both described by Monod functions and a fixed fraction of uptaken nutrients are invested into detoxification (Supplementary Methods). Placement of this system in a closed batch culture setting prevents allogenic influxes of intermediates and thus satisfies the purely autogenic assumption of the cEO equation. Calculation of the instantaneous intra-specific interaction 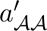 (Fig. 2D) recapitulates the environmental dependency of interactions in this system, with positive intra-specific interactions generally dominating at high toxin concentrations and negative intra-specific interactions dominating at low toxin concentrations. This static map is traversed by the system as it evolves from some initial state ***r***_0_, following the trajectory ***r***(*t*). We can see in Fig. 2E that in this particular case, the system generally moves towards the origin as *A* reduces the concentration of both the nutrient [*n*] and the toxin [*q*]. Importantly, this means that 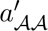 can switch from being positive early to being negative later on.

**Figure 2:**
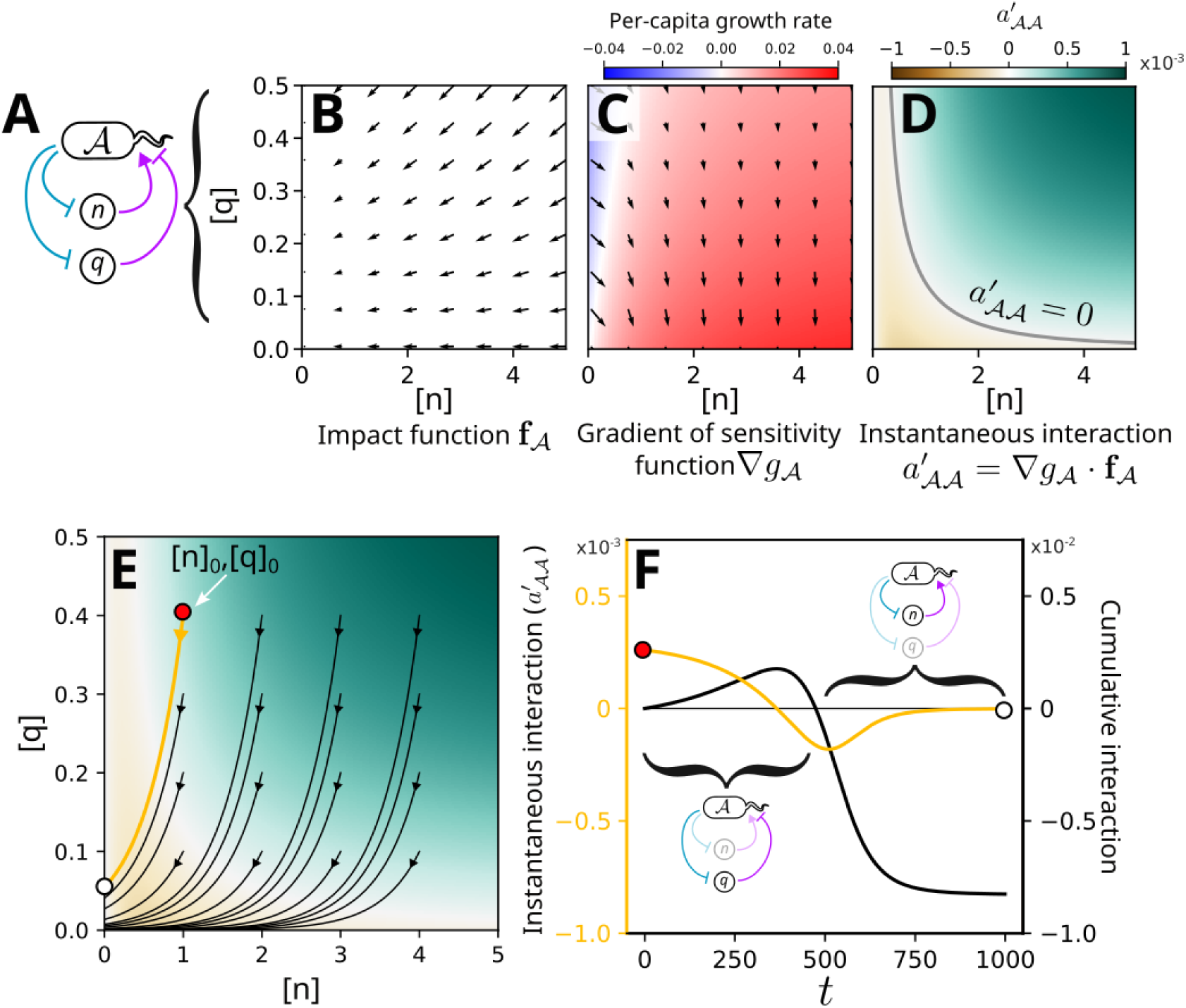
Intra-specific interactions mediated by mixtures of positive and negative mechanisms are predicted to switch sign over time in batch culture. **A** One of the simplest examples of a system with mixed pairwise elementary interactions is a single species *A* which increases the growth of other members of its population by detoxifying an environmental toxin while reducing their growth by depleting a common nutrient. **B**, **C** We can represent the impact and sensitivity functions for *A* using the ‘environment space’, which denotes the values of the different growth-limiting environmental factors (in this case, the concentrations of the nutrient [*n*] and of the toxin [*q*]). Impact functions are vector fields sitting in this space (black arrows, **B**), while sensitivity functions are scalar fields (**C**). The gradient of the sensitivity function then represents the direction in the environment space in which the growth rate of *A* increases most rapidly, as well as how quickly it increases (black arrows, **C**). **D** Taking the scalar product of the impact function and the gradient of the sensitivity function yields the instantaneous interaction 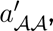 representing the instantaneous effect that *A* has on its own growth rate at a given position in the environment space. **E** Purely autogenic systems such as batch culture experiments trace out trajectories in this environment space, starting from an initial position [*n*]_0_, [*q*]_0_. **F** We can calculate both 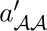 and the integrated effect of *A* on its own growth (main text), demonstrating a switch in the effective intra-specific interaction: at early timepoints, when the toxin concentration is high, detoxification dominates and the interaction appears positive. By contrast, at late timepoints when the toxin has mostly been removed, depletion of the single nutrient dominates and the instantaneous intra-specific interaction becomes negative.

This switch in sign of the instantaneous intra-specific interaction propagates through to *A*’s growth rate. As there are no other species in this system, the sole growth rate effect is the time-dependent impact of *A* on its own growth – the cumulative intra-specific interaction – given by 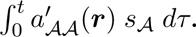 This switches from positive to negative once the accumulated benefit of the removal of the toxin is outweighed by the accumulated penalty from the reduction in the nutrient concentration (Fig. 2F). We therefore arrive at an unexpected prediction: measurements of the intra-specific interaction in such systems should give positive values if performed early on (when detoxification dominates) and negative values if performed later (when depletion dominates).

### 2.3 An antibiotic-based experimental system demonstrates sign-switching of the intra-specific interaction

We now attempted to establish whether this prediction was borne out in an experimental setting. As an experimental model of the detoxification/depletion interaction network (Fig. 2A), we made use of a bacterium (*Comamonas testosteroni*) that can degrade *β*-lactam antibiotics via induced secretion of *β*-lactamases (Fig. S1) and which can utilise proline as a sole carbon source. *β*-lactamase-producing bacteria are typically associated with a phenomenon known as the inoculum effect, in which larger starting population sizes result in higher measured Minimum Inhibitory Concentrations (MICs) of the antibiotic.^31,32^ This is due to more rapid degradation of the antibiotic at larger initial population sizes, an effect already suggestive of a positive intraspecific interaction. Combined with a depletion mechanism mediated by competition over limited proline as a sole carbon source, we speculated that we would observe a positive to negative intra-specific interaction shift in a time-dependent manner as predicted theoretically.

To address this, we prepared arrays of environmental conditions (with varying initial proline, [pro]_0_, and ampicillin, [amp]_0_, concentrations) within 96-well plates (Fig. 3A). Each condition was split into two sets of wells, one inoculated with exponential-phase *C. testosteroni* cells at high density and the second at low density. Absorbance-based growth curves of these cultures were then measured in a plate reader, which were used to calculate a quantity we call the *measured interaction* (for a comparison of the three interaction concepts we discuss in this paper – instantaneous, cumulative and measured – please refer to Supplementary Text 3 and Fig. S2).

**Figure 3:**
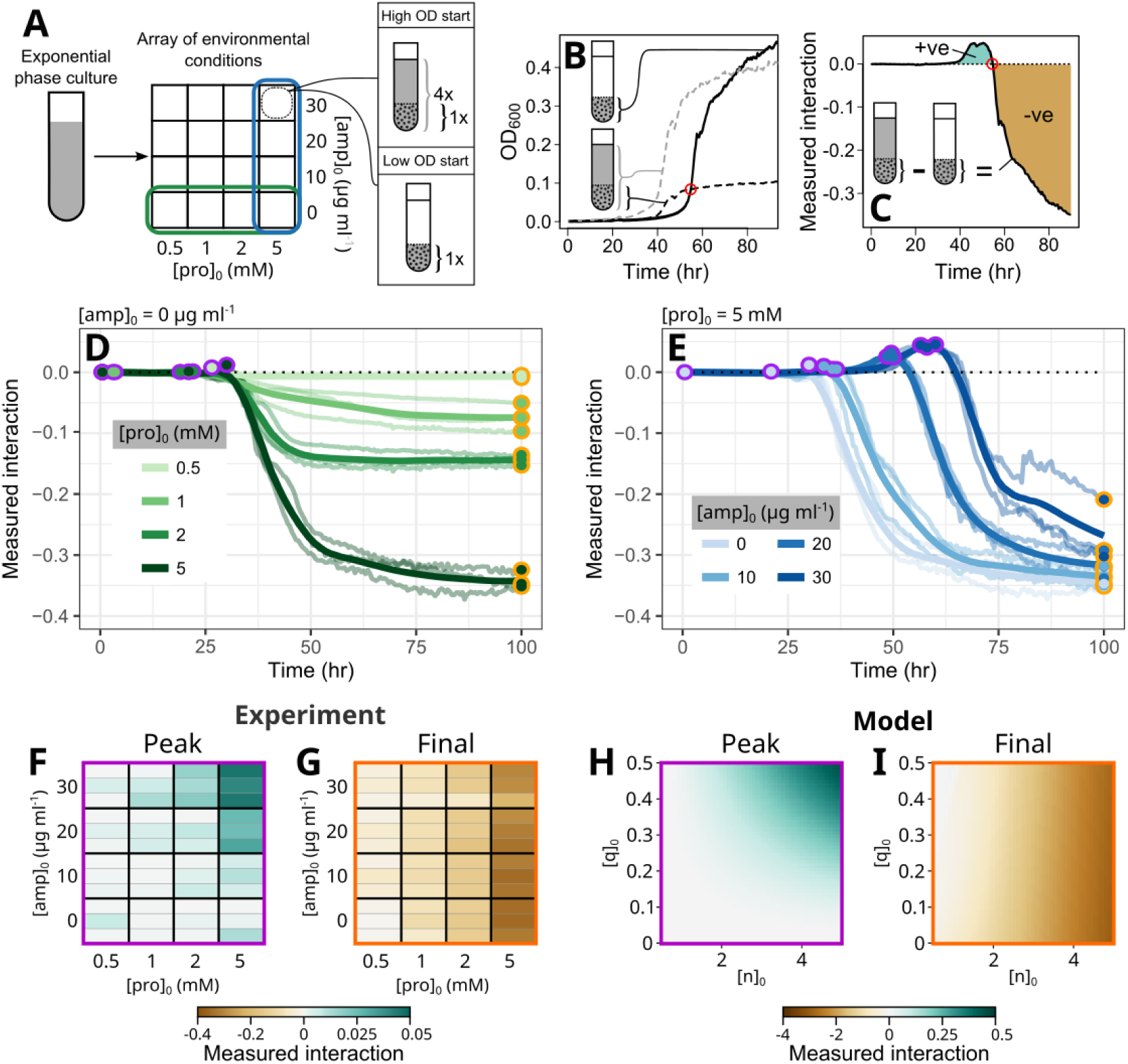
An antibiotic-based model system demonstrates sign switching of measured intra-specific interactions over time. *Comamonas testosteroni* is a *β*-lactamase producing soil bacterium which can degrade environmental ampicillin. Combined with competition over a single limiting carbon source (proline), we used this as an experimental analogue of the model shown in Fig. 2. **A** Exponential-phase cells were transferred to a 96-well plate containing wells with different initial ampicillin concentrations [amp]_0_ and proline concentrations [pro]_0_. Six wells were prepared for each condition, consisting of three replicates each of low and high initial inoculation densities at a 1:4 density ratio. **B**, **C** We measured the growth curve of each well and averaged the technical replicates. We then calculated the measured interaction over time by normalising the averaged high optical density (OD) curve by the ratio of the starting ODs (**B**) and subtracting the low OD curve (main text, Fig. S2, Supplementary Text 3) (**C**). Measured interactions greater than 0 indicate that growth of a matched sub-population of *C. testosteroni* (black dots) was enhanced by the presence of additional members of the same species in the high-OD wells relative to the low-OD wells (a positive intra-specific interaction), while differences less than 0 indicate growth suppression (a negative intra-specific interaction). **D**, **E** Comparing measured interactions across different proline (**D**) and ampicillin (**E**) concentrations demonstrates the environment-dependent shift in positive to negative interactions predicted by the model. We summarise this shift for each condition by measuring the peak (purple circles) and final (orange circles) measured interactions for each condition (**F**,**G**). **H**,**I** These qualitatively match predictions from our modelling framework. The general pattern that emerges from these simulations is robust to changes in simulation parameters (Fig. S4). Faint lines in **D** indicate *n* = 3 separate biological replicates performed on separate days, while bold lines indicate LOESS-smoothed averages. Biological replicates are indicated in **F** and **G** by separate horizontal strips.

Our definition of the measured interaction is based on typical experimental measurements of interactions, in which monoculture and co-culture assays are prepared with a constant inoculation density of a focal species. The interaction is then measured by detecting whether this initial population grows more or less in the presence of a second species.^9,33–35^ By direct analogy, we can treat our low inoculation density condition as a ‘monoculture-like’ assay, with a corresponding sub-population in the high-density condition which is of equal size. In the highdensity condition, this sub-population is effectively co-cultured with a second sub-population of the same species. We can therefore measure the intra-specific interaction by comparing the fate of the matching sub-populations in the high- and low-density conditions (Fig. 3B).

In practice, this is achieved by dividing the growth curve of the high-density culture by the ratio of inoculation densities (4:1), yielding the size of the sub-population as a function of time. At times when this normalised curve is higher than that of the low inoculation density condition, we can infer that the presence of additional cells of the same species enhanced the sub-population’s growth – *i.e.* that a positive intra-specific interaction has occurred. The opposite argument applies when the normalised curve is lower than that of the low inoculation density condition (Fig. 3B). To measure the intra-specific interaction as a function of time, we can therefore simply subtract the low inoculation density curve from the normalised high inoculation density curve (Fig. 3C). We considered several alternative definitions of the measured interaction (Fig. S2D-G), but found that the chosen abundance difference metric provided the optimal balance between capturing the shape of the cumulative interaction and robustness to measurement noise. We note that it is similar to accepted endpoint-based interaction metrics,^34^ although we emphasise that in contrast to these measurements which yield a single, fixed value, our approach yields a time-varying interaction estimate.

Beginning with the control conditions with zero antibiotic, we observe that the low inoculation density curves look very similar to the high inoculation density curves aside from a consistent time delay (Fig. S3A). We can interpret this delay as arising mostly from the smaller initial number of cells in the low density condition, as the ratio of densities between the two conditions remains approximately equal to the inoculation ratio until the high-density condition approaches stationary phase (at around 35 hrs). This is reflected in the measured intra-specific interaction, which is approximately neutral up to this point and negative afterwards (Fig. 3D). In the presence of antibiotic the time delay between the two conditions increases, presumably because the smaller initial population is slower to degrade the ampicillin before starting to grow (Fig. S3B). Consequently, we see a concentration-dependent positive interaction emerging with increasing ampicillin doses, as predicted by the model (Fig. 3E). Ultimately, all environments resulted in negative interactions in the long term. Summarising these experimental results by considering the peak and final measured interactions demonstrates the environmental and time dependencies together (Fig. 3F,G), which qualitatively match the patterns predicted by our modelling framework (Fig 3H,I). Although we do not directly fit model parameters to our data, we find that these qualitative patterns are robust to large changes in parameter values, suggesting that these results are not a result of fine-tuning of the model (Fig. S4).

Evolutionary rescue can result in similar abundance trajectories as those described here, as a small number of mutant cells with antibiotic-resistant genotypes can grow to fixation after a long time delay. ^36,37^ Although the consistency of our growth curves (Fig. S3) argued against evolutionary rescue, which typically leads to highly variable growth trajectories between replicates,^36^ we decided to test the role of evolution in our experimental system by measuring the MIC of ampicillin for each culture at the end of one of our interaction measurement timecourses (Fig. S5). These showed a small increase (*≈* 50%) in the resistance of populations exposed to the highest ampicillin concentrations compared to those grown under antibiotic-free conditions. However, simulations incorporating the evolution of resistance showed that evolutionary trends, far from driving the observed interaction time dependencies, tend to attenuate measured positive interactions if they have any effect at all (Fig. S6). Thus, we concluded that the consistent positive to negative interaction switch that we observe arises from the changing dominance of the two elementary interactions (detoxification and depletion), as suggested by our theoretical framework.

### 2.4 Small crossfeeding communities illustrate the common origins of time-dependent interactions and spatial structure

So far, we have considered time and environmental dependencies in a mono-species system. However, our framework generalises quite readily to multi-species communities, as well as certain types of spatially-structured communities (Methods, Supplementary Text 4, Fig. S7). Two recent studies have described time^15^ and spatial^38^ dependencies of very similar two-species communities. In both cases, a degrader species consumes a polymer (chitin or dextran) and subsequently produces a metabolite (acetate or glucose) which can be consumed by the second crossfeeding community member. When the polymer is exhausted, the degrader species can switch from net production to net consumption of the crossfed metabolite (Fig. 4A). Daniels et al. ^15^ note a time-dependency of the inter-specific interactions in batch culture: supply of an initial quantity of polymer leads initially to a strongly positive degrader *→* crossfeeder interaction and a neutral crossfeeder *→* degrader interaction, changing later to a weakly positive crossfeeder *→* degrader interaction and a weakly negative crossfeeder *→* degrader interaction (Fig. 4B). In a separate study, Wong et al. ^38^ loaded a similar community into a microfluidic device consisting of a single channel along which flow was applied. They observe that this community spontaneously self-structures along the channel, with the crossfeeding species only being able to grow towards its outlet (Fig. 4C). Given the commonalities between the two studies, we decided to use them as case studies for how our framework can unify similar observations occurring across time or space.

**Figure 4:**
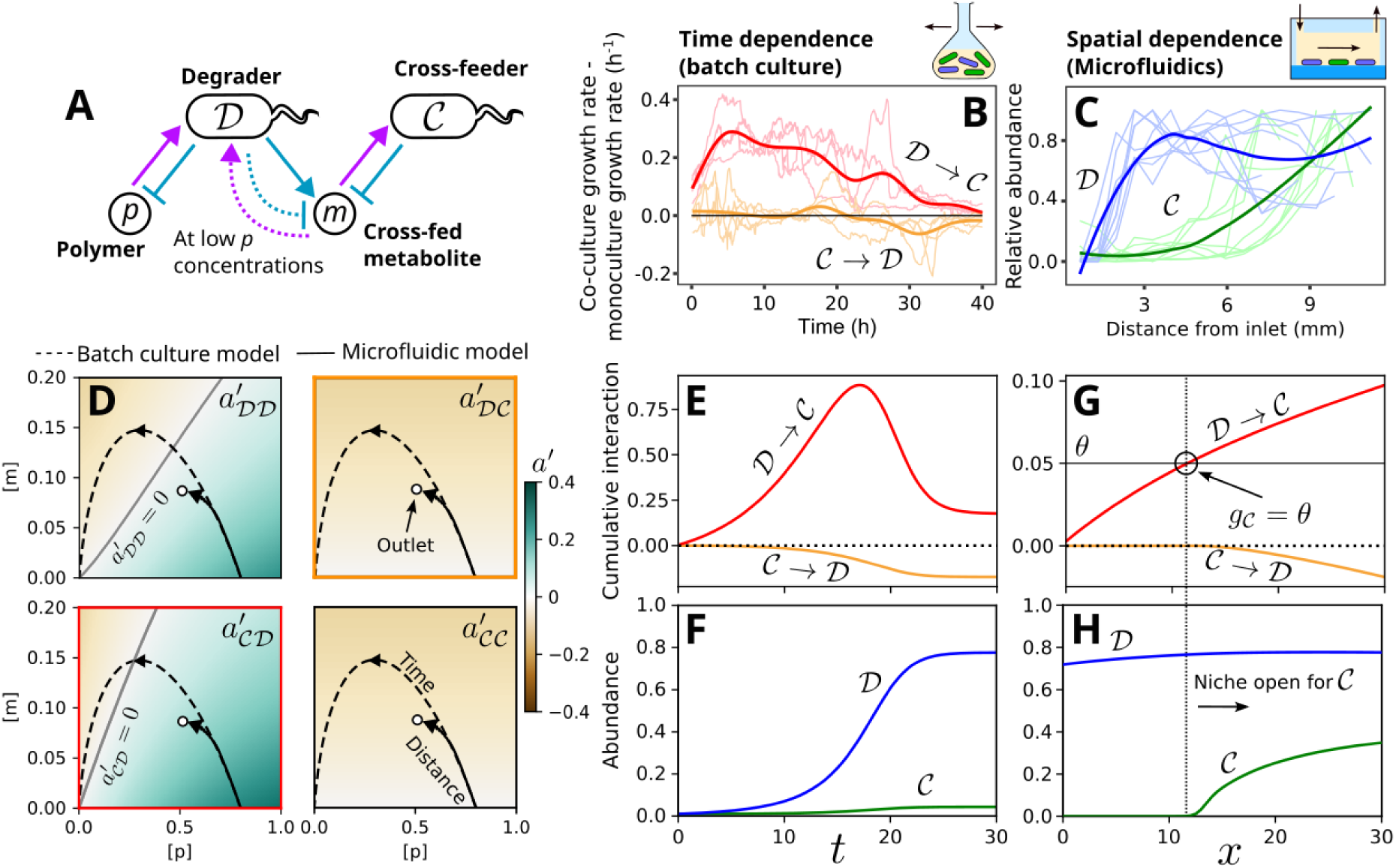
Our framework shows that interaction time dependencies and spatial structure can arise from closely-related processes. **A** Two recent studies^15,38^ describe the ecological patterns arising in a two-species community consisting of a degrader *D* that consumes a polymer *p* and produces a metabolic by-product *m* which is consumed by a second crossfeeding species (*C*). At low concentrations of *p*, *D* switches from net production of *m* to consumption. **B** Daniels et al. ^15^ find that this type of community displays time-dependent inter-specific interactions in batch culture, with the impact of *D* on *C* increasing early on and decreasing later (red) and the impact of *C* on *D* switching from neutral to negative (orange). **C** By contrast, Wong et al. ^38^ show how a similar community patterns itself in microfluidic channels with unidirectional flow, with *C* only being able to grow towards the outlet of the device. **D** We constructed an EO model of this community and applied our analytical techniques to obtain the instantaneous interaction matrix for each possible pair of community members (main text, Supplementary Methods). We then simulated the environmental trajectories of batch culture (dashed lines) and the microfluidic device (solid lines) inoculated with this community (Methods). In the case of the microfluidic device, the initial environment [*p*]_0_, [*m*]_0_ corresponds to the composition of the media injected into the system at the inlet, while points along the environmental trajectory indicate the steady-state media composition at different positions along the channel. **E**, **F** In the batch-culture model, the gradual enhancement of the environment by *D* for *C* via conversion of *p* to *m* results in a gradual increase in the cumulative interaction from *D* to *C*. Later, once *p* has been largely exhausted, the switch in the behaviour of *D* from net production to net uptake of *m* leads to competition between the two species, and a downward trend in both inter-specific cumulative interactions (**E**). These dynamics are difficult to dissect from the raw growth curves (**F**). **G**, **H** When this community is placed into the spatial context of a simulated microfluidic channel, we observe a similar interaction pattern from the inlet to the outlet, with a positive interaction accumulating from *D* to *C*. At a certain position, this positive cumulative interaction exceeds the mortality rate *θ* representing the flushing of cells by flow. Beyond this point, the net growth rate of *C* is positive, reflecting the opening of a niche for *C* (**G**). This leads to the spatial structuring of the two species observed in experiments (**H**).

We built an EO model of the degrader/crossfeeder community, labelling the degrader population *D* and the crossfeeder population *C*. We then derived expressions for the four different instantaneous interactions 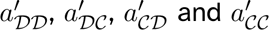 (Supplementary Methods). As shown in Fig. 4D, these four quantities can be arranged analogously to the interaction matrix of the gLV framework, with intra-specific interactions located along the main diagonal and inter-specific interactions located off this axis. However, instead of being represented by a single value as in the gLV approach, the instantaneous interactions are expanded into scalar fields defined on the entire environment space, capturing the environmental-dependency of each different interaction. Both the degrader’s intra-specific instantaneous interaction 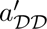 and inter-specific instantaneous interaction 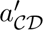 contain both positive and negative regions, reflecting the changing balance between the enrichment mechanism (production of the crossfed metabolite from the polymer) and the depletion mechanism (competition over the crossfed metabolite) in different environments.

In batch culture, organisms modify their environment by secreting and consuming intermediates over time. A similar effect occurs in flowing systems, whereby the intermediates within a parcel of fluid are sequentially modified by the organisms residing at successive spatial locations as it is transported downstream. We can therefore project trajectories representing the evolution of the environment over time in a batch culture system (Fig. 4D, dashed lines) and the spatial variation of the environment along the length of a system under flow at steady-state (Fig. 4D, solid lines) onto these instantaneous interaction maps, allowing us to interpret the changing interactions over time and space using the same framework (Methods, Supplementary Text 4). The initial position of the system ***r***_0_ – here assumed to consist of a large amount of polymer and zero crossfed metabolite – is interpreted subtly differently between the two cases: in batch culture, this represents the initial composition of the inoculum media, while in the microfluidic channel this represents the fixed composition of the media in the inflow of the device. Both systems sweep out initial trajectories with similar shapes, suggesting that the temporal patterning of the batch culture and the spatial patterning of the channel may arise from similar changes in interaction strengths.

To explore this in more detail, we now broke down the growth dynamics in the batch culture simulations into cumulative interactions, focusing on the inter-specific cases (Fig. 4E,F). We observe a similar pattern of the time evolution in the batch culture interactions as in the original study (Fig. 4B,E). In the initial phase, the large amount of initial polymer is metabolised by the degrader, resulting in large amounts of overspill in the form of the crossfed metabolite. This consequently substantially enhances the growth of the crossfeeder, while the relatively low utility of the crossfed metabolite at this point for the degrader limits the impact of its uptake by the crossfeeder on the growth of the degrader. Later, the switch of the degrader to metabolite uptake leads to a decrease in the strength of the net-positive interaction with the crossfeeder, and a negative impact of the crossfeeder on the growth of degrader.

Very similar effects arise as we consider the spatially structured system (Fig. 4G,H). Near the inlet, the crossfeeder cannot grow as the rate at which it is washed out of the device (*θ*) exceeds the growth rate sustained at very low metabolite concentrations. However, the activity of the degrader leads to a gradual enhancement of the environment for the crossfeeder along the length of the channel and ultimately leads to the opening of a new niche when the cumulative interaction from the degrader to the crossfeeder exceeds the threshold set by *θ*. This leads to the spatial structure observed in the original study, with the crossfeeder only growing towards the outlet of the device (Fig. 4C,H). Our model also reproduces the suppressive effect of increased flow rates on the growth of the crossfeeder, as observed experimentally: ^38^ higher flowrates cause the environmental trajectory to terminate with less monomer and polymer having been consumed due to more rapid wash-out of the two substrates (Fig. S8A). Combined with a higher mortality *θ* associated with the stronger flow, this ultimately prevents the crossfeeder’s niche from being efficiently opened up and halts its growth (Fig. S8B-G). In summary, our framework shows how spatial patterns arising under uni-directional flow and interaction time-dependencies in well-mixed systems are reflections of the same underlying ecological processes.

## 3 Discussion

We have presented a general framework that explains context dependencies of interactions as arising from feedbacks between organisms and their environment. This viewpoint provides a theoretical justification for the ubiquity of context dependencies of environmentally-mediated interactions:^13,39,40^ aside from some carefully chosen combinations of the impact and sensitivity functions, Eq. 4 implies that essentially *every* environmentally-mediated interaction will depend on the environmental state. Furthermore, as autogenic interaction mechanisms typically manifest through environmental change, interaction changes over time should be widespread. Our single-species toxin/nutrient system (Figs. 2, 3) provides an illustration of this effect. Initially the population interacts positively with itself (increases its own growth rate) through environmental detoxification, but this mechanism inherently causes an environmental change that results in a switch of the sign of the instantaneous interaction – once the toxin is eliminated, the positive interaction mechanism is suppressed and competition for the nutrient dominates. The autogenic environmental changes thus effectively set up a stress gradient in time, driving the observed time-dependency of the interaction.^25^

Our work also shows how placement of the same community of organisms in different types of system (*e.g.* batch culture *versus* chemostats *versus* flowcells) can result in distinct but connected phenomena. By revealing these underlying connections, we show how different experimental paradigms can be exploited to shed light on different aspects of the underlying ecology. For example, in Fig. 4 we show how time dependencies in batch culture and spatial structure in flowcells are manifestations of the same EO feedbacks. More practically, in Supplementary Texts 1 and 2 we discuss a novel route by which our framework allows one to predict interactions in balanced allogenic/autogenic systems by measuring interactions in fully autogenic experimental systems, thereby providing a basis for disentangling autogenic and allogenic contributions to interactions. Crucially, this approach is based purely on species abundance measurements, meaning it can be applied even when the underlying interaction mechanisms are unknown.

The cEO equation can be interpreted as a framework for predicting interaction changes, given a hypothesis about the underlying feedbacks between the organisms and their environment and the assumption that these autogenic feedbacks are the predominant mechanism driving interactions. Theoretical models that have attempted to capture context dependencies have generally done so via *ad hoc* manipulations of the interaction parameters. ^41,42^ Other studies explicitly take EO dynamics into account but stop short of drawing out the context-dependencies, leaving them implicitly encoded in the impact and sensitivity functions.^9,12,27,43–46^ Our approach shows how such dependencies emerge within such systems as a whole.

Our results have particular relevance for our understanding of the outcomes of batch culture interaction measurements. ^9,33–35,47,48^ The mechanism by which measured interactions in batch culture switch from positive to negative over time once a single nutrient becomes limiting (Fig. 3) is quite general, and suggests that measurements based on end-point abundances may miss positive interactions during early community establishment. This may at least partially explain the ongoing controversy surrounding the relative distribution of negative and positive interactions in natural communities, where the predominance of negative interactions as estimated by end-point batch culture methods appears to be at odds with findings based on alternative _approaches._^34,35,49–51^

We also note that despite our focus on microbial ecosystems, our results should also hold true for macroscopic ecosystems as long as the assumptions of our framework – particularly our assumption of autogenic dominance – are at least approximately true. Indeed, the role of the interplay between organisms and their environment has long been understood to drive primary succession in plant ecosystems, in which modification of the local environment by early pioneer species leads to the opening of new niches and eventual replacement of pioneers by latecomers better adapted for the new environment. ^52,53^ In Supplementary Text 5 and Fig. S9, we illustrate this idea by presenting a model of fully autogenic primary succession in a plant community. Decomposition of the changes in growth rates into separate cumulative interactions illustrates the complex time-dependency of the interactions in this system, with some interactions changing sign over the course of the succession (Fig. S9E). Similar successional patterns are observed in macroscopic systems such as whale falls^54^ and microscopic systems such as marine snow ^55,56^ in which allogenic nutrient fluxes are substantially smaller than the autogenic impacts of detritivores. Likewise, although we have focused on microfluidic channels as models of systems under unidirectional flow, analogous systems such as rivers and animal digestive tracts are widespread. The spatial niche-opening effects that our framework describes may therefore at least partially explain the longitudinal patterning of organisms in such systems.^57–59^

Nevertheless, there are some limitations to our framework. While we can generalise our framework to the case when allogenic mechanisms are present (Eq. S4), our unification of environmental, temporal and spatial patterns rests on the assumption that allogenic factors can be eliminated. This limiting assumption allows us to draw an analogy between the cEO and gLV equations, but runs contrary to communities in nature which are typically subject to external change. Nevertheless, this assumption is less limiting than it may appear at first glance, covering for example nutrient cycling in closed environments. ^60^ We also do not specify how the initial environment ***r***_0_ is reached prior to *t* = 0. In general, we envisage that some allogenic process sets this initial environmental state, such as the deposition of virgin substrate by a disturbance in the case of primary succession or the composition of an animal’s diet in the case of a longitudinally-structured gut microbiome. Ultimately however, the appropriate choice of ***r***_0_ is an external constraint which must be carefully selected by the modeller. Lastly, our assumption that interactions are environmentally-mediated, while well-grounded for many microbial and plant communities, ^52,61^ cannot account for direct interaction mechanisms such as predation and contact-dependent processes.^62,63^ These may be incorporated into Eq. 3 simply by adding the standard gLV description of density-dependent interactions (∑*_β_ a_αβ_s_β_*), although this hybrid interaction description is less simple to interpret than the base cEO equation.

In summary, our work shows that much of the diversity of interaction context-dependencies can be explained by reciprocal feedbacks between organisms and their environment. We give multiple examples of how explicit theoretical representation of these feedbacks can be used to predict and interpret interaction changes, providing a path forwards in the effort to manipulate interactions to predictable ends. Ultimately, we anticipate that a renewed focus on the dynamic role of the environment in determining the properties of ecosystems across scales will open new methods for controlling communities, as well as help to resolve longstanding questions regarding their composition and diversity.

## Acknowledgements

We would like to thank E. Ulrich, M. Amicone, S. Sulheim, C. Vulin, A. Del Panta, P.Padmanabha, G. Ugolini and J. Palmer for their valuable comments on a previous version of this manuscript. We would also like to thank J. Wong for sharing microfluidic data. Both authors were supported by the Swiss National Science Foundation (SNSF) through the NCCR Microbiomes (51NF40 180575), while OJM was additionally supported by a Human Frontier Science Program (HFSP) long-term fellowship (LT0020/2022-L) and SM was supported by an SNSF Eccellenza grant (PCEGP3 181272).

## 4 Data and code availability statement

All data and code used in this study (apart from data reproduced from other studies, Fig. 4B,C) is available at https://github.com/Mitri-lab/enviroInteracts.

## 5 Materials and methods

### Environment-Organisms models

Our toxin-nutrient model (Fig. 2A) was derived from a pre-existing framework, ^9^ while our degrader-crossfeeder model (Fig. 4A) was built from a Monod-based description of polymer/metabolite fluxes. We provide a complete description of these systems – including detailed derivations of instantaneous interactions – in the Supplementary Methods.

### Batch culture simulations

To generate trajectories representing the time evolution of batch culture experiments, the coupled systems of ODEs for both models were numerically integrated using scipy’s solve_ivp function, using the Runge-Kutta method of order 5(4). The starting population sizes for the toxin-nutrient model were 0.001 (low inocultion density) and 0.004 (high inoculation density), while both the degrader and crossfeeder populations were initiated at a density of 0.01 for the degrader-crossfeeder model. Initial environmental compositions (***r***_0_) are indicated in the relevant figures.

### Microfluidic simulations

Our simulations of spatially-structured flowing systems require that we move from a purely temporal model of intermediate and species changes to a spatio-temporal model. To do this, we represent the concentration profiles of the full set of intermediates as a set of 1D scalar fields ***r***(*x, t*), with the position along the channel represented by the spatial coordinate *x*. This varies from the inlet position *x* = 0 to the outlet position *x* = *L*. Likewise, the profile of the species abundances ***s*** is represented by the set of 1D scalar fields ***s***(*x, t*). Implicitly, we assume that the system is small enough in the *y* and *z* dimensions that it is effectively well-mixed along these axes by diffusion, allowing us to make use of the 1D approximation to study longitudinal structure.

We simulate the dynamics of the media composition using the 1D advection-diffusion equation:

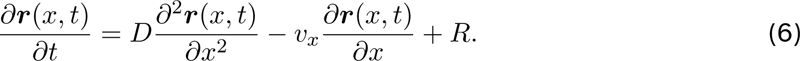

Here, on the right hand side, the first term represents the diffusive fluxes of intermediates along the length of the channel, with a rate set by the diffusion coefficient *D* = 0.5 which we take to be equal for all intermediates. This value is high enough to ensure numerical stability of the resulting environmental trajectories. The second term represents the advective fluxes mediated by active flow, with a rate set by the flow velocity *v_x_*. We choose values of *v_x_* to ensure that advection dominates over diffusion given the channel length *L* and the diffusion coefficient *D*, a necessary condition of our framework (Supplementary Text 4). The final term represents the sources and sinks of intermediates at each position, in this case given by an adjusted form of the impact functions for the degrader-crossfeeder model (Eq. S23). Together, these terms give the total rate of change of the intermediate concentrations at a particular location in the channel. Microbial population dynamics are simulated at each spatial location and are assumed to grow statically (*i.e.* to not be transported by diffusion or flow), with local dynamics based on an adjusted form of the sensitivity functions of the degrader-crossfeeder model based on the local concentrations of intermediates (Eq. S22). We describe the necessary adjustments to the impact and sensitivity functions in the Supplementary Methods, along with their implications for the calculation of the cumulative interaction.

To ensure the simulated composition of the inflowing media remains fixed, we set 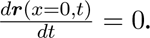 Additionally, we use an absorbing boundary condition at *x* = *L* to simulate free variation of the composition of the media at the outlet. The initial composition of the environment was set uniformly as ***r***(*x, t* = 0) = ***r***_0_, *i.e.* equal to the composition of the inflow. Both populations were seeded uniformly throughout the system at a density of ***s***(*x, t* = 0) = 0.01. The set of PDEs was numerically integrated using the solve_ivp function. We simulated dynamics for *t* = 1000 time units until the system reached a steady-state (Fig. S7), allowing application of our theoretical framework.

### Experiments

#### Strains and growth conditions

Our *C. testosteroni* strain MWF001 comes from a pre-existing study. ^9^ Cells were streaked onto TSA plates from freezer stocks and grown overnight. Single colonies were then picked (one colony per biological replicate), and cells grown overnight in glass Erlenmeyer flasks under continuous shaking in base minimal media (Table S1) supplemented with 10 mM proline. Due to the slow growth of *C. testosteroni* under these conditions, cells were in exponential phase at the end of this period. Cells were then washed twice in PBS. The OD_600_s of the washed cultures were then measured and cultures diluted to initialise experiments at the appropriate starting densities as described below. Cultures were grown at 28*^◦^*C in all cases.

#### Intra-specific interaction measurements

We prepared 96-well plates with a variety of environmental conditions by filling each well with 180 *µ*l of basal media supplemented with varying concentrations of proline ([pro]_0_ = 0.5, 1, 2, 5 mM) and ampicillin ([amp]_0_ = 0, 10, 20, 30 *µ*g ml*^−^*^1^). 20 *µ*l of an exponential-phase culture of *C. testosteroni* was then added, with three wells of each condition containing culture adjusted to high density (OD_600_ in well = 0.004) and three wells containing culture adjusted to low density (OD_600_ in well = 0.001). The plate was placed into a plate reader (BioTek Synergy H1) and OD_600_ readings for each well taken every 30 mins for 120 hr at 28 *^◦^*C under continual shaking between timepoints. The background signal was subtracted from the resulting raw growth curves by first estimating the OD contribution from the cells in the high inoculation OD wells (*κ*) using the equation

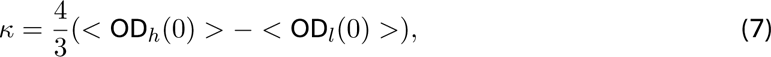

where *<* OD*_h_*(0) *>* and *<* OD*_l_*(0) *>* represent the plate-wide average initial OD readings for the high inoculation density and low inoculation density wells, respectively. The factor of 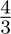 stems from the 1:4 inoculation density ratio. Each curve was now individually adjusted by subtracting the average OD of the specified curve’s first 3 timepoints and adding either *κ* for the high inoculation density wells or 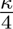 for the low inoculation density wells. The average OD curves were then calculated from the three replicates for each condition and used to calculate the measured interactions shown in Fig. 3D-G.

## References

[1] Pringle, E. G. PLOS Biology 2016, 14, e2000891.

[2] Lidicker, W. Z. BioScience 1979, 29, 475–477.

[3] Volterra, V. Nature 1926, 118, 558–560.

[4] Lotka, A. J. Proceedings of the National Academy of Sciences 1920, 6, 410–415.

[5] Novak, M.; Yeakel, J. D.; Noble, A. E.; Doak, D. F.; Emmerson, M.; Estes, J. A.; Jacob, U.; Tinker, M. T.; Wootton, J. T. Annual Review of Ecology, Evolution, and Systematics 2016, 47, 409–432.

[6] O’Dwyer, J. P. Theoretical Ecology 2018, 11, 441–452.

[7] MacArthur, R. Theoretical Population Biology 1970, 1, 1–11.

[8] Allesina, S.; Tang, S. Nature 2012, 483, 205–208.

[9] Piccardi, P.; Vessman, B.; Mitri, S. Proceedings of the National Academy of Sciences of the United States of America 2019, 116, 15979–15984.

[10] Hoek, T. A.; Axelrod, K.; Biancalani, T.; Yurtsev, E. A.; Liu, J.; Gore, J. PLOS Biology 2016, 14, e1002540.

[11] Martino, R. D.; Picot, A.; Mitri, S. PLOS Biology 2024, 22, e3002482.

[12] Rodríguez-Verdugo, A.; Vulin, C.; Ackermann, M. Ecology Letters 2019, 22, 838–846.

[13] Chamberlain, S. A.; Bronstein, J. L.; Rudgers, J. A. Ecology Letters 2014, 17, 881–890.

[14] Venkataram, S.; Kuo, H.-Y.; Hom, E. F. Y.; Kryazhimskiy, S. Nature Ecology and Evolution 2023, 7, 143–154.

[15] Daniels, M.; van Vliet, S.; Ackermann, M. The ISME Journal 2023 2023, 17, 1–11.

[16] Nadell, C. D.; Drescher, K.; Foster, K. R. Nature Reviews Microbiology 2016, 14, 589–600.

[17] Chang, C.-Y.; Bajić, D.; Vila, J. C. C.; Estrela, S.; Sanchez, A. Science 2023, 381, 343–348.

[18] Friedman, J.; Higgins, L. M.; Gore, J. Nature Ecology & Evolution 2017, 1, 1–7.

[19] Drew, G. C.; Stevens, E. J.; King, K. C. Nature Reviews Microbiology 2021, 19, 623–638.

[20] Liu, O. R.; Gaines, S. D. Proceedings of the National Academy of Sciences 2022, 119, e2118539119.

[21] Maron, J. L.; Baer, K. C.; Angert, A. L. Journal of Ecology 2014, 102, 1485–1496.

[22] Bertness, M. D.; Shumway, S. W. The American Naturalist 1993, 142, 718–724.

[23] Callaway, R. M.; Walker, L. R. Ecology 1997, 78, 1958–1965.

[24] Malkinson, D.; Tielbörger, K. Oikos 2010, 119, 1546–1552.

[25] Brooker, R. W.; Callaghan, T. V. Oikos 1998, 81, 196–207.

[26] Tilman, D. The American Naturalist 1980, 116, 362–393.

[27] Picot, A.; Shibasaki, S.; Meacock, O. J.; Mitri, S. Current Opinion in Microbiology 2023, 75, 102354.

[28] Meszéna, G.; Gyllenberg, M.; Pásztor, L.; Metz, J. A. Theoretical Population Biology 2006, 69, 68–87.

[29] Koffel, T.; Daufresne, T.; Klausmeier, C. A. Ecological Monographs 2021, 91, e01458.

[30] Estrela, S.; Libby, E.; Cleve, J. V.; Débarre, F.; Deforet, M.; Harcombe, W. R.; Peña, J.; Brown, S. P.; Hochberg, M. E. Trends in Ecology and Evolution 2019, 34, 6–18.

[31] Parker, R. F. Experimental Biology and Medicine 1946, 63, 443–446.

[32] Lenhard, J. R.; Bulman, Z. P. Journal of Antimicrobial Chemotherapy 2019, 74, 2825–2843.

[33] Mitri, S.; Richard Foster, K. Annual Review of Genetics 2013, 47, 247–273, PMID: 24016192.

[34] Foster, K. R.; Bell, T. Current Biology 2012, 22, 1845–1850.

[35] Kehe, J.; Ortiz, A.; Kulesa, A.; Gore, J.; Blainey, P. C.; Friedman, J. Science Advances 2021, 7, 7159.

[36] Orr, H. A.; Unckless, R. L. PLOS Genetics 2014, 10, e1004551.

[37] Ramsayer, J.; Kaltz, O.; Hochberg, M. E. Evolutionary Applications 2013, 6, 608–616.

[38] Wong, J. P. H.; Fischer-Stettler, M.; Zeeman, S. C.; Battin, T. J.; Persat, A. Proceedings of the National Academy of Sciences of the United States of America 2023, 120, e2217577120.

[39] He, Q.; Bertness, M. D.; Altieri, A. H. Ecology Letters 2013, 16, 695–706.

[40] Shantz, A. A.; Lemoine, N. P.; Burkepile, D. E. Ecology Letters 2016, 19, 20–28.

[41] Hernandez, M. J. Proceedings of the Royal Society B: Biological Sciences 1998, 265, 1433– 1440.

[42] Holland, J. N.; Deangelis, D. L. Ecology Letters 2009, 12, 1357–1366.

[43] Goldford, J. E.; Lu, N.; Bajić, D.; Estrela, S.; Tikhonov, M.; Sanchez-Gorostiaga, A.; Segrè, D.; Mehta, P.; Sanchez, A. Science 2018, 361, 469–474.

[44] Müller, M. J. I.; Neugeboren, B. I.; Nelson, D. R.; Murray, A. W. Proceedings of the National Academy of Sciences of the United States of America 2014, 111, 1037–1042.

[45] Hammarlund, S. P.; Chacón, J. M.; Harcombe, W. R. Environmental Microbiology 2019, 21, 759–771.

[46] Momeni, B.; Xie, L.; Shou, W. eLife 2017, 6.

[47] Hsu, R. H.; Clark, R. L.; Tan, J. W.; Ahn, J. C.; Gupta, S.; Romero, P. A.; Venturelli, O. S. Cell Systems 2019, 9, 229–242.e4.

[48] Weiss, A. S. et al. The ISME Journal 2021 16:4 2021, 16, 1095–1109.

[49] Palmer, J. D.; Foster, K. R. Science 2022, 376, 581–582.

[50] Yu, J. S. et al. Nature Microbiology 2022, 7, 542–555.

[51] Zelezniak, A.; Andrejev, S.; Ponomarova, O.; Mende, D. R.; Bork, P.; Patil, K. R. Proceedings of the National Academy of Sciences of the United States of America 2015, 112, 6449–6454.

[52] Roberts, D. W. Vegetatio 1987, 69, 27–33.

[53] Connell, J. H.; Slatyer, R. O. The American Naturalist 1977, 111, 1119–1144.

[54] Smith, C. R.; Glover, A. G.; Treude, T.; Higgs, N. D.; Amon, D. J. Annual Review of Marine Science 2015, 7, 571–596.

[55] Pontrelli, S.; Szabo, R.; Pollak, S.; Schwartzman, J.; Ledezma-Tejeida, D.; Cordero, O. X.; Sauer, U. Science Advances 2022, 8, eabk3076.

[56] Datta, M. S.; Sliwerska, E.; Gore, J.; Polz, M. F.; Cordero, O. X. Nature Communications 2016, 7, 11965.

[57] Vannote, R. L.; Minshall, G. W.; Cummins, K. W.; Sedell, J. R.; Cushing, C. E. Canadian Journal of Fisheries and Aquatic Sciences 1980, 37, 130–137.

[58] Riva, A.; Kuzyk, O.; Forsberg, E.; Siuzdak, G.; Pfann, C.; Herbold, C.; Daims, H.; Loy, A.; Warth, B.; Berry, D. Nature Communications 2019, 10, 1–11.

[59] Pereira, F. C.; Berry, D. Environmental Microbiology 2017, 19, 1366–1378.

[60] de Jesús Astacio, L. M.; Prabhakara, K. H.; Li, Z.; Mickalide, H.; Kuehn, S. Proceedings of the National Academy of Sciences 2021, 118, e2013564118.

[61] Gralka, M.; Szabo, R.; Stocker, R.; Cordero, O. X. Current Biology 2020, 30, R1176–R1188.

[62] Sockett, R. E. Annual Review of Microbiology 2009, 63, 523–539.

[63] Hayes, C. S.; Aoki, S. K.; Low, D. A. Annual Review of Genetics 2010, 44, 71–90.

