## Supplementary Information for "Environment-organism feedbacks drive changes in ecological interactions"

### Environment-organism feedbacks drive changes in ecological interactions – Supplementary Information

#### 1 Supplementary Text

##### Supplementary Text 1: Derivation of the cEO, eEO and fEO equations

To obtain our expression for the per-capita growth rate of a species $\alpha$ in a purely autogenic Environment-Organism system (Eq. 3), we begin from our definitions of the impact and sensitivity functions (Eqs. 2 and 1). We assume that the system sweeps out some trajectory $\Gamma$ in the environment space which can be parameterized by the variable $\tau$. $\tau$ can be interpreted as representing the history of the system up to the current query time $t$. At $\tau = 0$, the system begins at an initial position $\boldsymbol{r}_0$ in the environment space, while at $\tau = t$ it has reached an end position $\boldsymbol{r}(t)$. We seek to express the per-capita growth rate at this query time.

By our definition of the sensitivity function, we have

$$\frac{1}{s_\alpha}\frac{ds_\alpha}{dt} = g_\alpha(\boldsymbol{r}). \tag{S.1}$$

From the gradient theorem, we can rewrite this as

$$g_\alpha(\boldsymbol{r}) = g_\alpha(\boldsymbol{r}_0) + \int_\Gamma \nabla g_\alpha(\boldsymbol{r}) \cdot d\boldsymbol{r}, \tag{S.2}$$

where the second term is the path integral of the gradient of the sensitivity function over the trajectory $\Gamma$. We can use the definition of a path integral over a vector field to express this as

$$\begin{aligned}\int_\Gamma \nabla g_\alpha(\boldsymbol{r}) \cdot d\boldsymbol{r} &= \int_0^t \nabla g_\alpha(\boldsymbol{r}) \cdot \frac{d\boldsymbol{r}}{d\tau} d\tau \\ &= \int_0^t \nabla g_\alpha(\boldsymbol{r}) \cdot \left( \boldsymbol{\sigma} + \sum_\beta \boldsymbol{f}_\beta(\boldsymbol{r})\ s_\beta \right)\ d\tau \\ &= \int_0^t \nabla g_\alpha(\boldsymbol{r}) \cdot \boldsymbol{\sigma}\ d\tau + \sum_\beta \int_0^t \nabla g_\alpha(\boldsymbol{r}) \cdot \boldsymbol{f}_\beta(\boldsymbol{r})\ s_\beta\ d\tau.\end{aligned} \tag{S.3}$$

We find the term $\nabla g_\alpha(\boldsymbol{r}) \cdot \boldsymbol{f}_\beta(\boldsymbol{r})$ naturally arising in this expression. Calling this $a'_{\alpha\beta}(\boldsymbol{r})$, we can put together a general expression for ecosystems for which the allogenic environmental influence $\boldsymbol{\sigma}$ can vary over time. We will call this the ‘fluctuating Environment-Organism’ (fEO) equation:

$$\frac{1}{s_\alpha}\frac{ds_\alpha}{dt} = g_\alpha(\boldsymbol{r}_0) + \int_0^t \nabla g_\alpha(\boldsymbol{r}) \cdot \boldsymbol{\sigma}\ d\tau + \sum_\beta \int_0^t a'_{\alpha\beta}(\boldsymbol{r})\ s_\beta\ d\tau. \tag{S.4}$$

For purely allogenic systems, we can assume that $\boldsymbol{\sigma} = \boldsymbol{0}$, thus we obtain the ‘closed Environment-Organism’ (cEO) equation:

$$\frac{1}{s_\alpha}\frac{ds_\alpha}{dt} = g_\alpha(\boldsymbol{r}_0) + \sum_\beta \int_0^t a'_{\alpha\beta}(\boldsymbol{r})\ s_\beta\ d\tau. \tag{S.5}$$

Systems with a constant allogenic input $\boldsymbol{\sigma} \neq \boldsymbol{0}$ – such as chemostats – can reach a long-term equilibrium. To describe this, we can set $\frac{1}{s_\alpha}\frac{ds_\alpha}{dt} = 0$ in Eq. S.4 and assume that a steady-state environmental composition $\boldsymbol{r}^*$ is reached. Noting that $\sum_\beta \int_0^t a'_{\alpha\beta}(\boldsymbol{r})\ s_\beta\ d\tau$ must be equal and opposite to $\int_0^t \nabla g_\alpha(\boldsymbol{r}) \cdot \boldsymbol{\sigma}\ d\tau$ for all times $t$ to satisfy the equilibrium condition, we can write down the ‘equilibrium Environment-Organism’ (eEO) equation:

$$-\nabla g_\alpha(\boldsymbol{r}^*) \cdot \boldsymbol{\sigma} = \sum_\beta a'_{\alpha\beta}(\boldsymbol{r}^*)\ s_\beta, \tag{S.6}$$

which is exactly equivalent to the gLV equation at equilibrium, taking $\nabla g_\alpha(\boldsymbol{r}^*) \cdot \boldsymbol{\sigma}$ as the intrinsic growth rate and $a'_{\alpha\beta}(\boldsymbol{r}^*)$ as the elements of the interaction matrix.[1] This equation describes a situation where allogenic and autogenic influences on organismal growth are exactly balanced. An equivalent expression has previously been discussed in Meszena et al.[2] and Koffel et al.,[3] though we provide a new derivation of this result.

#### Supplementary Text 2: Measuring instantaneous interactions in novel systems

In most of this paper, we consider systems in which we know the underlying EO dynamics, represented by our knowledge of the impact and sensitivity functions. However, when investigating a novel community this information is generally not available. It would therefore be useful to find a way of measuring instantaneous interactions purely based on observable data, specifically population abundances over time.

The typical approach to measuring interactions in microbiology is to compare the outcomes of monoculture and co-culture experiments.[4,5] However, as we have emphasised throughout this work, EO interactions depend strongly on environmental context. Two factors thus make such comparisons difficult: environmental trajectories are generally different in monoculture and co-culture settings, and the environmental state is typically unobservable at a given time. We therefore face the problem of finding times in monoculture and co-culture datasets at which the environmental composition $\boldsymbol{r}$ is equal, which are the times at which co-culture/monoculture comparisons would be valid.

Fortunately, there is one time at which the environmental composition can be forced to be equal between co-culture and monoculture conditions – at the beginning of the experiment, when the environment is set at $\boldsymbol{r}_0$. We can exploit this fact to measure $a'_{\alpha\beta}$ around $\boldsymbol{r}_0$.

We begin with the cEO equation (Eq. S.5):

$$\frac{1}{s_\alpha}\frac{ds_\alpha}{dt} = g_\alpha(\boldsymbol{r}_0) + \sum_\beta \int_0^t a'_{\alpha\beta}(\boldsymbol{r})\ s_\beta\ d\tau. \tag{S.7}$$

If we consider a sufficiently short time window at the beginning of a batch culture experiment, the environment at the end of this window $\boldsymbol{r}_t$ will be close to the initial environment $\boldsymbol{r}_0$ and so the instantaneous interaction will be approximately constant. Thus we obtain the approximation:

$$g_\alpha(\boldsymbol{r}_t) \approx g_\alpha(\boldsymbol{r}_0) + \sum_\beta a'_{\alpha\beta}(\boldsymbol{r}_0) \int_0^t s_\beta\ d\tau. \tag{S.8}$$

For a monoculture batch culture experiment of species $\alpha$, we have:

$$g_\alpha(\boldsymbol{r}_t) \approx g_\alpha(\boldsymbol{r}_0) + a'_{\alpha\alpha}(\boldsymbol{r}_0) \int_0^t s_\alpha \, d\tau, \tag{S.9}$$

thus the initial instantaneous intra-specific interaction can be approximated as:

$$a'_{\alpha\alpha}(\boldsymbol{r}_0) \approx \frac{g_\alpha(\boldsymbol{r}_t) - g_\alpha(\boldsymbol{r}_0)}{\int_0^t s_\alpha \, d\tau}. \tag{S.10}$$

We can now consider a co-culture between $\alpha$ and a partner $\beta$:

$$g_\alpha(\boldsymbol{r}_t) \approx g_\alpha(\boldsymbol{r}_0) + a'_{\alpha\alpha}(\boldsymbol{r}_0) \int_0^t s_\alpha \, d\tau + a'_{\alpha\beta}(\boldsymbol{r}_0) \int_0^t s_\beta \, d\tau, \tag{S.11}$$

which can be rearranged to give the initial instantaneous inter-specific interaction between $\beta$ and $\alpha$ in terms of the abundances of the two populations and the intra-specific instantaneous interaction obtained from monoculture measurements:

$$a'_{\alpha\beta}(\boldsymbol{r}_0) \approx \frac{g_\alpha(\boldsymbol{r}_t) - g_\alpha(\boldsymbol{r}_0) - a'_{\alpha\alpha}(\boldsymbol{r}_0) \int_0^t s_\alpha \, d\tau}{\int_0^t s_\beta \, d\tau}. \tag{S.12}$$

These equations therefore allow us to approximate the entire 'instantaneous interaction matrix' $a'_{\alpha\beta}(\boldsymbol{r}_0)$ at $\boldsymbol{r}_0$ using only monoculture and co-culture growth rate data. Environmental context-dependencies can then be quantified by measuring how interactions change between different starting environments.

These measurements are rather limited, in that they only allow $a'_{\alpha\beta}$ to be measured at a single location in the environment space ($\boldsymbol{r}_0$). However, if this location is carefully chosen, we can obtain very useful information about communities that can be used to assess their long-term stability and feasibility. In particular, if $\boldsymbol{r}_0$ is set as $\boldsymbol{r}^*$ – the steady-state environmental composition for an ecosystem at autogenic/allogenic equilibrium (Eq. S.6) – this procedure allows us to obtain the interaction matrix in the autogenic/allogenic setting based on measurements from purely autogenic settings (Supplementary Text 1). In a non-microbial context, we might carry out this procedure by cutting off an environment at equilibrium at $\boldsymbol{r}^*$ from all allogenic influences (for example, by transplanting soil at a location of interest to controlled greenhouse conditions) and growing pairs of species within that enclosed system.

We note lastly that the early per-capita growthrates $(g_\alpha(\boldsymbol{r}))$ required by these expressions are generally strongly influenced by measurement noise when extracted from growth curves, as very small early population sizes result in a low signal to noise ratio. This noise is further inflated when the population-level growth rate is divided by the small population size to yield the per-capita growth rate $(g_\alpha(\boldsymbol{r}) = \frac{1}{s_\alpha}\frac{ds_\alpha}{dt})$. A similar effect can be observed in Fig. S2D,E, which led to our rejection of similar interaction measurement candidates in the main text. To avoid this, one can use a late cutoff time $t$ that allows populations to grow to sizes that are large relative to the measurement noise, but this risks violating the assumption that $\boldsymbol{r}_t \approx \boldsymbol{r}_0$. However, recent methodological advances allow single-cell growth rates in temporally varying environments to be measured directly via microscopy,[6] potentially allowing this problem to be circumvented in the future.

#### Supplementary Text 3: Glossary of interaction definitions

In the main text, we discuss three different definitions of 'interaction' for EO systems: instantaneous interactions, cumulative interactions and measured interactions. For convenience, we summarise this hierarchy of terms here, highlighting the relationships between them.

- **Instantaneous interaction:** The per-capita rate at which species $\beta$ is impacting the growth rate of species $\alpha$ through its modification of environmental factors. Using the EO formulation, this is given as the scalar product of the impact function of $\beta$ and the gradient of $\alpha$'s sensitivity function at a given point in the environment space $\boldsymbol{r}$ ($a'_{\alpha\beta}(\boldsymbol{r}) = \nabla g_{\alpha}(\boldsymbol{r}) \cdot \boldsymbol{f}_{\beta}(\boldsymbol{r})$). The instantaneous interaction is a second-order derivative with respect to time (*i.e.* an acceleration-like term), distinguishing it from the typical elements of an interaction matrix which are first-order derivatives.[1]

- **Cumulative interaction:** The total impact that species $\beta$ has had on the growth rate of $\alpha$ up to time $t$ through its modification of the environment. In the EO formulation this is given as $\int_0^t a'_{\alpha\beta}(\boldsymbol{r})\ s_{\beta}\ d\tau$. Like the elements of the interaction matrix this is a first-order derivative with respect to time, but unlike the interaction matrix it is not expressed in per-capita terms.

- **Measured interaction:** The total impact the presence of species $\beta$ on the abundance of $\alpha$ at time $t$, comparing between monoculture and pairwise co-culture experiments.[7] As the environmental trajectories $\boldsymbol{r}(t)$ will be different between monoculture and co-culture conditions there is no simple expression for this in terms of instantaneous or cumulative interactions, but denoting the abundance of $\alpha$ in monoculture as $s_{\alpha}^{m}(t)$ and in co-culture as $s_{\alpha}^{c}(t)$ – themselves given by integration of the cEO equation under monoculture and co-culture starting conditions (Eq. S.5) – the measured interaction is given simply as $s_{\alpha}^{c}(t) - s_{\alpha}^{m}(t)$.

In the case of intra-specific interactions (Figs. 2,3, S2), we inoculate high-density batch cultures at a starting density that is a factor of $R$ times larger than that of low-density batch cultures. Treating the additional members of the focal species $\alpha$ in this high-density culture as the co-cultured population $\beta$, the sub-population that is matched to the low-density culture is given by dividing through the growth curve for the high-density culture by $R$ (Fig. 2B). Thus, taking the abundance in the low-density culture as $s_{\alpha}^{l}(t)$ and in the high-density culture as $s_\alpha^h(t)$, the measured interaction is given as $\frac{s_\alpha^h(t)}{R} - s_\alpha^l(t)$.

This is a less natural definition than the instantaneous and cumulative interactions, being chosen to facilitate measurement of interactions from noisy experimental data (Fig. S2D-G). However, it fulfils several important requirements of an interaction measure. Firstly, it returns a value of zero when populations are not interacting: under non-limiting conditions, a population grows exponentially at a rate $\gamma$. Population sizes at time $t$ under high-density and low-density conditions are then given as $s_\alpha^h(t) = s_0 e^{\gamma t}$ and $s_\alpha^l(t) = \frac{s_0}{R} e^{\gamma t}$, respectively, where $s_0$ is the inoculation density of the high-density culture. The measured interaction is then given as $\frac{s_\alpha^h(t)}{R} - s_\alpha^l(t) = 0$. Secondly, measurements of long-term competition are insensitive to inoculation density. In batch culture, limiting nutrients are typically converted to biomass with a fixed yield (Fig. S5A), thus for a given initial quantity of nutrients a total of $G$ cells will be added over the growth period. The measured interaction at large $t$ is then given as $\frac{s_\alpha^h(t \to \infty)}{R} - s_\alpha^l(t \to \infty) = \frac{s_0 + G}{R} - \left(\frac{s_0}{R} + G\right) = G\left(\frac{1}{R} - 1\right)$, in which the inoculation density $s_0$ is eliminated. A corollary of this result is that the measured interaction will always be negative, corresponding to the assumption of pure nutrient competition in this example.

#### Supplementary Text 4: Use of the cEO framework to model systems under flow

In using our cEO framework to simulate the dynamics of systems subject to flow, we need to be careful to specify the conditions under which our assumption of autogenic dominance ($\boldsymbol{\sigma} = 0$) holds true. Indeed, by having an inlet into which fresh nutrients can be continuously fed, we appear to be grossly violating the assumption of zero allogenic input. However, we can consider a fluid parcel travelling along the length of the system to behave as a purely autogenic system (in the sense that changes to its composition are only mediated by the local microbial populations at position $x$) as long as the rate of flow is large relative to the diffusion rate of intermediates.

We can show this formally by beginning with the 1D advection-diffusion equation used to specify the evolution of the distribution of intermediates along the system over time (Eq. 6):

$$\frac{\partial \boldsymbol{r}(x,t)}{\partial t} = D\frac{\partial^2 \boldsymbol{r}(x,t)}{\partial x^2} - v_x \frac{\partial \boldsymbol{r}(x,t)}{\partial x} + R. \tag{S.13}$$

We proceed by making two assumptions: firstly, we assume that the system reaches a steady-state concentration profile $\boldsymbol{r}^*(x)$ (associated with a static distribution of species $\boldsymbol{s}^*(x)$), allowing us to set this equation to zero. Secondly, we assume that the Péclet number $Pe$, which is defined as the ratio of the advective to diffusive transport rates, is substantially greater than 1. This allows us to neglect the diffusive term in the advection-diffusion equation. We therefore obtain

$$v_x \frac{d\boldsymbol{r}^*(x)}{dx} = R = \sum_{\beta} s^*_{\beta}(x) \boldsymbol{f}_{\beta}(\boldsymbol{r}^*(x)). \tag{S.14}$$

We note the similarity of this expression to our original definition of the rate of change of the environment in a batch culture system (Eq. 1). We can match these two expressions exactly by noting that $v_x = \frac{dx}{dt}$ and reparameterising the environmental trajectory $\boldsymbol{r}^*(x)$ in terms of the time coordinate of a fluid parcel relative to the time of its emergence at the inlet ($x = 0$). We can use this same reparameterisation to write Eq. 2 in terms of the steady-state concentration profile as traversed by the fluid parcel. These can finally be combined together to obtain a spatial version of the cEO equation, and a corresponding definition of the cumulative interaction (Supplementary Methods). Thus, as long as we can justify the two original assumptions, our framework should be directly applicable to such systems.

Beginning with the assumption that the system approaches a stationary distribution, we show in Fig. S7 the environmental trajectories traced out along the length of the channel as a function of increasing simulated time. We observe, as expected, convergence on a static distribution $\boldsymbol{r}^*(t)$ over long timescales. It is difficult to define precise criteria under which this convergence will occur for general EO models, but a necessary condition is that the sensitivity functions $g_\alpha$ must contain a negative mortality term against which positive growth rates can be balanced in the long term. In our model, this is provided by $\theta$, which describes the rate at which cells are washed off the surface of the channel by flow. An additional factor which we have found to be important is the inclusion of a density-dependent mechanism for capping growth at a particular spatial location, given in our case by the capacity $\lambda$. In the absence of this, species that are able to grow on the source media simply accumulate indefinitely at the inlet, rather than spreading out along the channel. Addition of these elements requires an adjustment to how the cumulative interaction is calculated, which we specify in the Supplementary Methods.

Now turning to $Pe$, we note that it is defined as $Pe = \frac{Lv_x}{D}$, where $L$ is the characteristic length-scale of the system (in this case, the length of the channel). We can readily obtain approximate values for the diffusion constants of dextran and glucose (approximately $3 \times 10^{-8}$ cm$^2$ s$^{-1}$ and $5 \times 10^{-6}$ cm$^2$ s$^{-1}$, respectively). Furthermore, in the experiments described in Wong et al.,[8] $L = 2$ cm and $v_x$ varied between $0.0003$ cm s$^{-1}$ and $0.003$ cm s$^{-1}$. This gives us a range of $Pe$ between 120 and 200,000, both substantially higher than 1 and firmly placing these experiments in the advection-dominated regime. We have also chosen the equivalent model parameters such that $Pe$ varies between 75 and 1200, similarly placing the model in the advection-dominated regime.

#### Supplementary Text 5: Context-dependence of interactions during primary succession of plant communities

We mainly focus in this manuscript on microbial communities due to extent of control we have over their environment. However, our framework can be used to describe other types of ecosystem with similar properties. For example, plants predominantly interact with each other through indirect, environmentally-mediated mechanisms such as nutrient competition and shading.[9,10] Here, we develop an EO model describing primary succession in a simple plant community, showing how our approach can be used to untangle the complex dynamics of changes in interactions during succession.

The predominant mechanistic basis for primary succession has been a subject of debate for over a century.[11–13] Rather than reopen this discussion, we will state our two main assumptions, both of which have some support in the experimental literature and are likely to be applicable in at least some settings. Firstly, we assume that environmental change during the successional window is mostly driven by autogenic mechanisms (*e.g.* accumulation of soil, stabilisation of substrate) rather than by allogenic factors (*e.g.* weathering of substrate, climactic variations).[14,15] Elimination of allogenic factors is equivalent to setting the external input term $\sigma$ of Eq. S.4 to zero. Secondly, we assume that late successional species are principally excluded from early environments because local environmental conditions prevent their growth. This niche-based view is contrasted to an arrival-based succession mechanism, in which the life history traits of pioneers enable them to arrive at newly exposed sites earlier and grow to maturity quicker than latecomers.[12,16] These assumptions allow us to make use of the cEO equation (Eq. S.5) to untangle the changing roles of competition and facilitation during primary succession.

The impact and sensitivity functions for this system are heavily inspired by the resource ratio model of Tilman,[17] which assumes an investment trade-off between foliage and nutrient fixation. Plants therefore exist on a continuum between shade-intolerant/soil-enriching species to shade-tolerant/soil-depleting species (Fig. S9A). We initialise an equal quantity of each member of this community in an environment with high light availability and low soil quality, representing environmental conditions in virgin terrain such as glacier forelands[18] or volcanic landscapes.[15,19] Considering the species abundances over time (Fig. S9B), we see the classic successional pattern of shade intolerant, hardy plants giving way to increasingly shade tolerant, soil-dependent species over time, showing that the assumptions of our framework are compatible with successional dynamics.

Our main departure from Tilman[17] is how environmental change arises. Tilman's approach is an equilibrium-based one: the supply of light and soil is assumed to be set by an external allogenic process at a timescale that is sufficiently long for the local community to equilibrate before the next environmental change occurs. The sequence of equilibrium species abundances supported by the sequence of environmental compositions – chosen externally by the modeller – is then treated as the successional pattern. By contrast, the autogenic changes to the environment that drive the succession in our model (Fig. S9C,D) are fundamentally non-equilibrium phenomena that arise out of the underlying EO dynamics. Interestingly, the environmental trajectory is not a straightforward march towards high-quality soil and low light availability as previously assumed;[17] soil quality declines during the transition from species $1$ to $2$ as the latter species quickly takes up the nutrients deposited into the soil by $1$.

We can now use our interaction framework to disentangle the contributions of each species to this successional pattern (Fig. S9E). Several patterns emerge. Firstly, the initial pioneer species $1$ consistently improves the environment for the later species in the successional sequence but is weakly inhibited by them once they establish, leading to its eventual replacement by these later species. Likewise, the final species $4$ makes the environment worse for all other species as well as itself, preventing invasion by earlier populations and ultimately stabilising its own population. The dynamics for the two species forming the middle of this sequence ($2$ and $3$) are more complex. In general, they worsen environmental conditions for other populations as they establish and sequester light and nutrients. Evidence for this inhibitory effect is well-documented, with earlier species tending to suppress the seedlings of later species.[18,20] However, longer-term enrichment of the soil gradually improves environmental conditions until later species in the succession can establish.[15,18,19] These later populations then outcompete the earlier species by overshading them, driving them to extinction and further enriching the soil upon their decomposition. This is clearest for the net effect of species $\beta = 2$ on $\alpha = 4$, which switches sign from negative to positive over the timecourse. Thus, the net effect of one species on another can change from competitive to facilitative depending on the point in the successional sequence. Some evidence for such a switch in the interaction sign over time has previously been reported,[19] although in general the extremely long timescales of primary succession has hampered longitudinal studies of interaction changes.

In summary, this example shows how interaction values can change in complex ways during successional dynamics, demonstrating sign switching of interaction values depending on whether a population is growing (reducing environmental quality) or declining (increasing environmental quality). Our framework helps to tease apart these complex effects, explaining how a changing balance between competitive and facilitative interaction mechanisms over time may determine the overall dynamics of successional change.

### 2 Supplementary methods

#### Modelling

##### Toxin-nutrient model

Our single-species toxin-nutrient model (Fig. 2A) is adapted from a previously described EO framework.[21] We model the growth rate of a single species $\mathcal{A}$, with abundance denoted as $s_\mathcal{A}$, as being positively dependent upon the concentration of a nutrient $[n]$ and negatively dependent upon the concentration of a toxin $[q]$. We assume that both of these positive and negative growth impacts are saturating functions of their respective intermediates, modelled as Monod functions. We further assume that the concentrations of the intermediates are measured in units of the half-velocity constant of the two Monod terms and that a fraction $f$ of the utilised nutrient is invested into detoxification and the remainder into growth. Together, these assumptions give the per-capita growth rate as

$$\frac{1}{s_\mathcal{A}}\frac{ds_\mathcal{A}}{dt} = (1-f)\frac{\nu_n[n]}{[n]+1} - \frac{\nu_q[q]}{[q]+1}, \tag{S.15}$$

where $\nu_n$ is the maximal nutrient uptake rate and $\nu_q$ is the maximal toxin impact.

The impact functions of the original model incorporate a yield parameter that describes the efficiency at which resources are converted into biomass.[21] However, we can remove this parameter by non-dimensionalisation if we define the bacterial abundance $s_\mathcal{A}$ as being measured in terms of the amount of biomass produced by one unit of nutrient with zero toxin degradation investment ($f = 0$) and zero toxin ($[q] = 0$). Then we can write the nutrient dynamics as

$$\frac{d[n]}{dt} = -s_\mathcal{A}\frac{\nu_n[n]}{[n]+1}. \tag{S.16}$$

Toxin dynamics are modelled similarly but involve an additional term, the detoxification efficiency $\delta$, which sets the amount of toxin removed from the environment for each unit of nutrient invested. Thus we obtain

$$\frac{d[q]}{dt} = -s_\mathcal{A}\delta f[q]\frac{\nu_n[n]}{[n]+1}. \tag{S.17}$$

For simplicity, we do not incorporate a passive (nutrient-independent) toxin uptake term as in.[21]

Defining the environment vector $\boldsymbol{r} = \binom{[n]}{[q]}$, the impact function $\boldsymbol{f}_{\mathcal{A}}(\boldsymbol{r})$ can be written from Eqs. S.16 and S.17 as

$$\boldsymbol{f}_{\mathcal{A}}(\boldsymbol{r}) = -\frac{\nu_n[n]}{[n]+1}\begin{pmatrix}1\\ \delta f[q]\end{pmatrix}. \tag{S.18}$$

We note that Eq. S.15 is already in the form of a sensitivity function $g_{\mathcal{A}}(\boldsymbol{r})$. We can therefore derive its gradient as

$$\nabla g_{\mathcal{A}}(\boldsymbol{r}) = \begin{pmatrix}\frac{(1-f)\nu_n}{([n]+1)^2}\\ \frac{-\nu_q}{([q]+1)^2}\end{pmatrix}, \tag{S.19}$$

and so,

$$a'_{\mathcal{A}\mathcal{A}} = \nabla g_{\mathcal{A}}(\boldsymbol{r}) \cdot \boldsymbol{f}_{\mathcal{A}}(\boldsymbol{r}) = \frac{\nu_n[n]}{[n]+1}\left(\nu_q\delta f\frac{[q]}{([q]+1)^2} - (1-f)\frac{\nu_n}{([n]+1)^2}\right), \tag{S.20}$$

with a non-trivial nullcline $a'_{\mathcal{A}\mathcal{A}} = 0$ at

$$[n] = -1 + \sqrt{\frac{(1-f)\nu_n([q]+1)^2}{[q]f\delta\nu_q}}. \tag{S.21}$$

Parameter values for this model as used in this manuscript are given in table 2. These were not explicitly fitted to our experimental data but were chosen to qualitatively match our experimental results, representing a regime in which the toxin has a comparable negative growth rate impact as the nutrient's positive impact ($\nu_n = \nu_q$) but is efficiently degraded with a relatively low degradation investment. Given our non-dimensionalisation of the half-velocity constants, the range of initial concentrations $[n]_0$ and $[q]_0$ shown in Fig. 3H,I is also of significance; our choice of $0.5 < [n]_0 < 5$ and $[q]_0 < 0.5$ indicates that the nutrients are close to saturation for most of the simulated conditions, while the toxin will have an approximately linearly concentration-dependent effect. Large changes to these parameter values do not generally have a strong effect on the overall interaction patterns, although we do observe qualitatively different outcomes when the strength of the toxin compared to the nutrient is increased to such an extent that it entirely abolishes growth in either the low inoculation density condition or both conditions (Fig. S4).

##### Degrader-crossfeeder model

We model the system described in Fig. 4A by considering the dynamics of the degrader population $\mathcal{D}$, the crossfeeder population $\mathcal{C}$, the polymer $p$ and the crossfed metabolite $m$. We assume that $\mathcal{D}$ consumes $p$ according to Monod kinetics, utilising a fixed fraction $1 - \phi$ for its own growth and converting the remaining fraction $\phi$ into $m$. This is in turn utilised by both $\mathcal{C}$ and $\mathcal{D}$ for growth, again according to Monod kinetics.

We note that our model is structured such that $\mathcal{D}$ has priority access to the breakdown products of $p$. In similar systems, these breakdown products act as extracellular public goods, with both degraders and crossfeeders having equal access to them.[22] However, in the studies we consider here, two similar mechanisms preserve the priority status of the degrader: in Wong et al.,[8] the degrader *Bacteroides thetaiotaomicron* appears to import dextran and degrade it internally, as suggested by the upregulation of various nutrient importers and cytosolic dextranases in the presence of dextran. The breakdown products - principally glucose - can therefore largely be maintained as internal private goods, with the excess leaking out and acting as the nutrient source for the crossfeeder *Bacteroides fragilis*. Similarly, while the degrader *Vibrio natriegens* of Daniels et al.[6] does appear to release digestive enzymes to break down chitin externally, the resulting breakdown products cannot be metabolised directly by the crossfeeder *Alteromonas macleodii*. Instead, they must first be converted to acetate by the internal catabolic metabolism of the degrader. Thus, the crossfeeder must again wait until the substrate has passed through a stage in which it is a private good of the degrader, effectively separating the two species into different trophic levels.[23]

Denoting the environment vector $\boldsymbol{r} = \begin{pmatrix} [p] \\ [m] \end{pmatrix}$ and subscripting yield parameters $Y$, half-saturation constants $K$ and maximal rate constants $\nu$ with labels denoting species and intermediate, this model is described by the sensitivity functions

$$g_{\mathcal{D}}(\boldsymbol{r}) = (1-\phi)\frac{\nu_{\mathcal{D}p}[p]}{[p]+1} + \frac{\nu_{\mathcal{D}m}[m]}{[m]+K_{\mathcal{D}m}}, \tag{S.22a}$$

$$g_{\mathcal{C}}(\boldsymbol{r}) = \frac{\nu_{\mathcal{C}m}[m]}{[m]+K_{\mathcal{C}m}}. \tag{S.22b}$$

Note that a similar non-dimensionalisation has been applied here as for the toxin-nutrient model, with the polymer concentration being measured in terms of the half-saturation constant for the degrader ($K_{\mathcal{D}p}$).

We also have corresponding impact functions given by:

$$\boldsymbol{f}_{\mathcal{D}}(\boldsymbol{r}) = \begin{pmatrix} -\frac{\nu_{\mathcal{D}p}[p]}{[p]+1} \\ \phi\frac{\nu_{\mathcal{D}p}[p]}{[p]+1} - \frac{1}{Y_{\mathcal{D}m}}\frac{\nu_{\mathcal{D}m}[m]}{[m]+K_{\mathcal{D}m}} \end{pmatrix}, \tag{S.23a}$$

$$\boldsymbol{f}_{\mathcal{C}}(\boldsymbol{r}) = \begin{pmatrix} 0 \\ -\frac{1}{Y_{\mathcal{C}m}}\frac{\nu_{\mathcal{C}m}[m]}{[m]+K_{\mathcal{C}m}} \end{pmatrix}, \tag{S.23b}$$

where the yield constant $Y_{\mathcal{D}p}$ has likewise been eliminated via non-dimensionalisation, along with a factor specifying the number of units of metabolite produced per unit of polymer.

The gradients of the sensitivity functions are given as

$$\nabla g_{\mathcal{D}}(\boldsymbol{r}) = \begin{pmatrix} (1-\phi)\frac{\nu_{\mathcal{D}p}}{([p]+1)^2} \\ \frac{\nu_{\mathcal{D}m}K_{\mathcal{D}m}}{([m]+K_{\mathcal{D}m})^2} \end{pmatrix}, \tag{S.24a}$$

$$\nabla g_{\mathcal{C}}(\boldsymbol{r}) = \begin{pmatrix} 0 \\ \frac{\nu_{\mathcal{C}m}K_{\mathcal{C}m}}{([m]+K_{\mathcal{C}m})^2} \end{pmatrix}, \tag{S.24b}$$

leading to the instantaneous interactions

$$a'_{\mathcal{CC}}(\boldsymbol{r}) = -\frac{1}{Y_{\mathcal{C}m}} \frac{\nu_{\mathcal{C}m}^2 K_{\mathcal{C}m}[m]}{([m]+K_{\mathcal{C}m})^3}, \tag{S.25a}$$

$$a'_{\mathcal{CD}}(\boldsymbol{r}) = \frac{\nu_{\mathcal{C}m} K_{\mathcal{C}m}}{([m]+K_{\mathcal{C}m})^2} \left( \phi \frac{\nu_{\mathcal{D}p}[p]}{[p]+1} - \frac{1}{Y_{\mathcal{D}m}} \frac{\nu_{\mathcal{D}m}[m]}{[m]+K_{\mathcal{D}m}} \right), \tag{S.25b}$$

$$a'_{\mathcal{DC}}(\boldsymbol{r}) = -\frac{1}{Y_{\mathcal{C}m}} \frac{\nu_{\mathcal{D}m} K_{\mathcal{D}m}}{([m]+K_{\mathcal{D}m})^2} \frac{\nu_{\mathcal{C}m}[m]}{([m]+K_{\mathcal{C}m})}, \tag{S.25c}$$

$$a'_{\mathcal{DD}}(\boldsymbol{r}) = (\phi - 1) \frac{\nu_{\mathcal{D}p}^2[p]}{([p]+1)^3} + \frac{\nu_{\mathcal{D}m} K_{\mathcal{D}m}}{([m]+K_{\mathcal{D}m})^2} \left( \phi \frac{\nu_{\mathcal{D}p}[p]}{[p]+1} - \frac{1}{Y_{\mathcal{D}m}} \frac{\nu_{\mathcal{D}m}[m]}{[m]+K_{\mathcal{D}m}} \right). \tag{S.25d}$$

The non-trivial nullcline of the inter-specific instantaneous interaction $a'_{\mathcal{CD}} = 0$ is given by

$$[p] = \frac{[m]\nu_{\mathcal{D}m}}{\phi \nu_{\mathcal{D}p} Y_{\mathcal{D}m}([m]+K_{\mathcal{D}m}) - \nu_{\mathcal{D}m}[m]}. \tag{S.26}$$

The nullcline of the intra-specific instantaneous interaction $a'_{\mathcal{DD}} = 0$ was found numerically using scipy's root function.

Parameter values for this model are given in table 3. Similarly to the toxin-nutrient model these are not explicitly fitted to the experimental data, and indeed we expect that they are substantially different between the two systems used in the studies we discuss here.[6,8] Such a qualitative approach does, however, allow us to determine the minimal differences between the underlying metabolic processes needed to bring about the observed interaction patterns in both systems. In particular, we find that we can reproduce the observed patterns if all metabolic processes (polymer degradation by the degrader and monomer uptake by both species) have equivalent kinetics ($\nu_{\mathcal{D}p} = \nu_{\mathcal{D}m} = \nu_{\mathcal{C}m}$ and $K_{\mathcal{D}m} = K_{\mathcal{C}m}$), the yields of the monomer and polymer are equal ($Y_{\mathcal{D}m} = Y_{\mathcal{C}m}$) and a fairly high proportion of polymer is converted to monomer and secreted ($\phi = 0.6$). We also assume that the polymer is introduced at a concentration below the corresponding half-velocity constant of the degrader ($[p]_0$=0.8), meaning the rate of polymer degradation will be strongly impacted by concentration changes over time and/or space.

##### Primary succesion model

Our model of primary succession in plant communities (Fig. S9) is heavily inspired by Tilman's formulation of the resource-ratio hypothesis.[17] We assume that plants compete for two resources, light arriving at the soil with irradiance $[l]$ and soil nutrients with concentration $[s]$, arranged in the environment vector $\boldsymbol{r} = \begin{pmatrix} [l] \\ [s] \end{pmatrix}$. Resource uptake and growth are modelled assuming Monod kinetics and overall rate limitation by the most limiting resource (Liebig's law of the minimum). For a given species $\mathcal{X}$ ($\mathcal{X} \in 1, 2, 3, 4$), the growth rate is then given as:

$$g_{\mathcal{X}}(\boldsymbol{r}) = \nu_{\mathcal{X}} \min\left(\frac{[l]}{[l] + K_{\mathcal{X}l}}, \frac{[s]}{[s] + K_{\mathcal{X}s}}\right) - \theta_{\mathcal{X}}. \tag{S.27}$$

For simplicity, we assume that all species have an equal maximal growth rate $\nu_{\mathcal{X}}$ and mortality rate $\theta_{\mathcal{X}}$. We further assume a trade-off between the half-velocity constants $K_{\mathcal{X}l}$ and $K_{\mathcal{X}s}$, representing different light versus nutrient acquisition strategies between different species;[17] specifically, we assume that $K_{\mathcal{X}s} K_{\mathcal{X}l} = 0.5$ (Table 4).

The gradient of this sensitivity function is then:

$$\nabla g_{\mathcal{X}}(\boldsymbol{r}) = \begin{cases} \begin{pmatrix} 0 \\ \nu_{\mathcal{X}} \frac{K_{\mathcal{X}s}}{([s]+K_{\mathcal{X}s})^2} \end{pmatrix} & \text{if } \frac{[s]}{[s]+K_{\mathcal{X}s}} < \frac{[l]}{[l]+K_{\mathcal{X}l}}, \\ \begin{pmatrix} \nu_{\mathcal{X}} \frac{K_{\mathcal{X}l}}{([l]+K_{\mathcal{X}l})^2} \\ 0 \end{pmatrix} & \text{if } \frac{[s]}{[s]+K_{\mathcal{X}s}} > \frac{[l]}{[l]+K_{\mathcal{X}l}}. \end{cases} \tag{S.28}$$

The impact functions for the effects of plants light and soil quality reflect the indirect mechanisms of shading and competition for soil nutrients through which plants interact.[9] Each new unit of species $\mathcal{X}$ reduces light availability by a fraction $\zeta_{\mathcal{X}}$ and increases it by an equivalent fraction upon its death. A population of $Y_{g\mathcal{X}}$ is yielded upon depletion of one unit of soil quality, and soil quality is increased upon the death and decay of one unit of $\mathcal{X}$ by $Y_{d\mathcal{X}}$; the product $Y_{\mathcal{X}d} Y_{\mathcal{X}g}$ thus determines the net enrichment of the soil mediated by the full life cycle of $\mathcal{X}$, taking into account fixation of nutrients such as nitrogen and bulking of soil volume with organic matter. Both $Y_{d\mathcal{X}}$ and $\zeta_{\mathcal{X}}$ are assumed to vary according to the sequence of the soil/light trade-offs, with light-intensive plants shading less and improving soil quality more than shade-tolerant species (Table 4). Explicitly, the impact function for $\mathcal{X}$ is given as:

$$\boldsymbol{f}_{\mathcal{X}}(\boldsymbol{r}) = \begin{pmatrix} -\zeta_{\mathcal{X}} \frac{[l]}{l_{\max}} \left(\nu_{\mathcal{X}} \min\left(\frac{[l]}{[l]+K_{\mathcal{X}l}}, \frac{[s]}{[s]+K_{\mathcal{X}s}}\right) - \theta_{\mathcal{X}}\right) \\ -\frac{1}{Y_{g\mathcal{X}}} \nu_{\mathcal{X}} \min\left(\frac{[l]}{[l]+K_{\mathcal{X}l}}, \frac{[s]}{[s]+K_{\mathcal{X}s}}\right) + Y_{d\mathcal{X}} \theta_{\mathcal{X}} \end{pmatrix}, \tag{S.29}$$

where $l_{\text{max}}$ is the maximal light availability with no foliage.

In Fig. S9, we initialise this system at $\boldsymbol{r} = \binom{3}{0.02}$, representing a starting state with high light availability and low soil quality. Species are seeded at the initial timepoint at a uniform density of 0.005.

#### Adjustments to cEO framework in microfluidic model

As we discuss in Supplementary Text 2, application of our framework to systems under flow requires that the dynamics approach a steady-state. To ensure this, we make two additions to our impact and sensitivity functions. Firstly, we assume that the microbial populations can grow only to a maximal density at a given location along the device, given by the channel capacity $\lambda = 1$. This is applied to each population independently in order to prevent a inter-specific density dependence, which would act as a direct interaction which could not be integrated into our framework; the maximal *total* abundance at a given site is therefore equal to $\lambda$ multiplied by the number of different populations. Both growth rates and associated impacts on concentrations of intermediates slow down as this capacity is approached. Secondly, we assume that microbes are flushed out of the system by flow at a rate $\theta = 0.005\ v_x$, analogous to the wash-out term in chemostat models.[24] The dependence of this wash-out rate on the flow rate $v_x$ is based on observations that suggest that biofilms are more strongly eroded at higher flow rates.[8,25] Microbial growth rates are therefore given as

$$\frac{1}{s_\alpha(x,t)}\frac{ds_\alpha(x,t)}{dt} = \frac{\lambda - s_\alpha(x,t)}{\lambda} g_\alpha(\boldsymbol{r}(x,t)) - \theta, \tag{S.30}$$

while the effective impact rates on the intermediates are given as

$$\frac{d\boldsymbol{r}(x,t)}{dt} = \sum_\beta \boldsymbol{f}_\beta(\boldsymbol{r}(x,t)) s_\beta(x,t) \frac{\lambda - s_\beta(x,t)}{\lambda}. \tag{S.31}$$

This latter expression is the explicit form of the term $R$ in Eq. 6.

The introduction of this density-dependent scaling term in the dynamics of the intermediates also requires an adjustment in the way the cumulative interaction is calculated. This arises because this term slows down the simulated metabolic rate of cells at high densities, reducing their effective impact on their environment. Once steady-state is achieved (Supplementary Text

2), the microbial abundances and resource concentrations become dependent solely on the position in the channel ($\boldsymbol{r}(x,t) \rightarrow \boldsymbol{r}^*(x)$ and $\boldsymbol{s}(x,t) \rightarrow \boldsymbol{s}^*(x)$). The cumulative interaction from $\beta$ to $\alpha$ is therefore given as $\frac{1}{v_x}\int_0^x a'_{\alpha\beta}(\boldsymbol{r}^*)s^*_\beta \frac{\lambda - s^*_\beta}{\lambda} d\chi$. Here, $\chi$ acts as a variable that parameterises the spatial trajectory of the system from the inlet to the query position $x$, analogously to how $\tau$ parameterises the temporal trajectory of batch-culture systems in Eq. 3.

#### Experimental methods

##### $\beta$-lactamase activity measurements

Exponential-phase cultures of *C. testosteroni* were inoculated into two Erlenmeyer flasks containing 20 ml of minimal media supplemented with 5mM proline at a starting $OD_{600}$ adjusted to 0.00025. To one of these flasks we added ampicillin at a label concentration of 100 $\mu$g ml$^{-1}$, however subsequent experiments suggested that degradation of our antibiotic freezer stocks had reduced the effective concentration to $\approx$ 30 $\mu$g ml$^{-1}$. The resulting cultures were grown under continuous shaking, and samples taken at 0, 6, 30, 54 and 78 hr. To measure the extracellular $\beta$-lactamase activity of cultures, samples were first spun down and the supernatant pipetted off. The enzymatic activity of the supernatant was then measured using a $\beta$-lactamase activity assay kit (Sigma-Aldrich, MAK221). In Fig. S1, the $\beta$-lactamase detection limit is defined as two times the standard deviation of the signal estimated from sample-free control wells.

##### MIC measurements

To determine the change in the resistance of *C. testosteroni* to ampicillin over the duration of the intra-specific interaction assay, we measured the Minimum Inhibitory Concentration (MIC) of ampicillin for each of the cultures in the 96-well plate at the end of one biological replicate of our experiment. Concentration gradients of ampicillin were first prepared in 96-well plates containing LB media, to which we added samples from the interaction measurement plate (specifically, from the low inoculation density wells as these were subject to the strongest selective pressure). As the $\beta$-lactamase resistance mechanism results in different MICs based on the starting density of culture (the inoculum effect[26,27]), we adjusted the inoculation volume to ensure approximately equal numbers of cells were added regardless of the final density of the cultures being measured. To do this, we took advantage of the fact that the final density of the samples was directly proportional to the initial proline concentration $[\text{pro}]_0$ (Fig. S5A), allowing us to simply scale the inoculation volume by $[\text{pro}]_0$. Specifically, we used 20 $\mu$l of $[\text{pro}]_0$ = 0.5 mM cultures, 10 $\mu$l of $[\text{pro}]_0$ = 1 mM cultures, 5 $\mu$l of $[\text{pro}]_0$ = 2 mM cultures and 2 $\mu$l of $[\text{pro}]_0$ = 5 mM cultures. These inoculua resulted in starting densities around 10 times greater than those of the interaction assay, explaining why the resulting MICs were substantially higher than the values of $[\text{amp}]_0$ used during interaction measurements. The total volume in each well was fixed at 200 $\mu$l. Plates were incubated at 28°C under continuous shaking. Final MICs were defined as the lowest concentration of ampicillin at which the $OD_{600}$ of a given sample was reduced by at least 80% relative to an antibiotic-free control after 20 hrs.

### 3 Supplementary figures

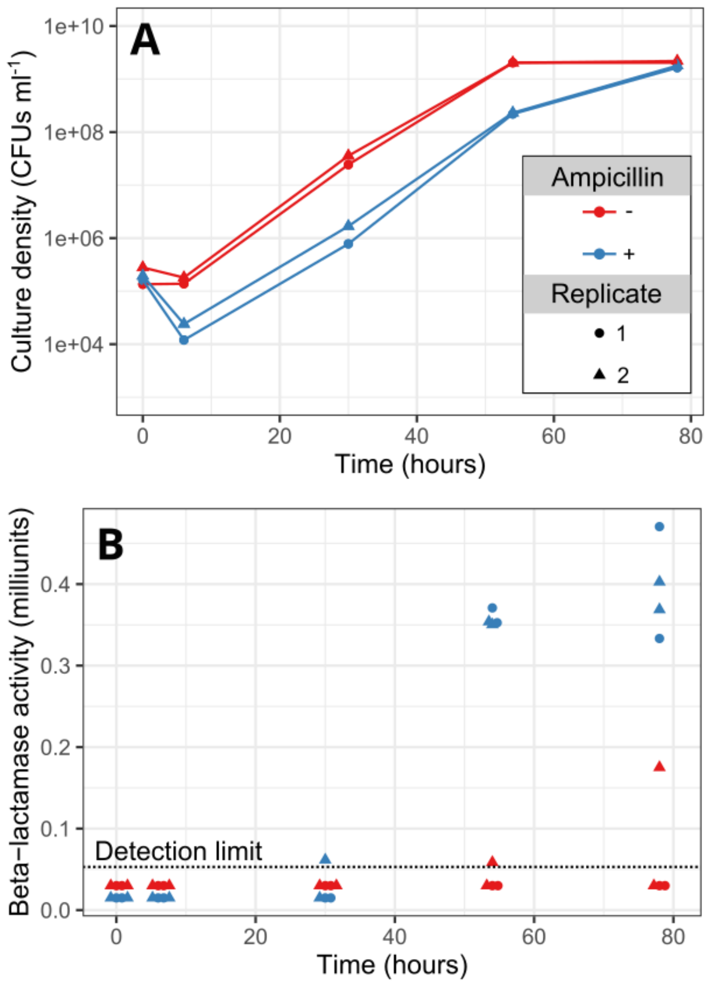


Figure S1: ***C. testosteroni* secretes $\beta$-lactamases when grown with ampicillin.** $n = 2$ cultures of *C. testosteroni* initialised from separate colonies were grown in media containing 5 mM proline and either with (blue) or without (red) ampicillin. Samples were taken from cultures at the indicated timepoints and cell densities (**A**) and $\beta$-lactamase activities (**B**) measured. One unit of $\beta$-lactamase is defined as the amount of enzyme needed to hydrolyse 1 nmol of nitrocefin in 1 ml of solution in 1 minute. Points with equal shapes and colours in **B** indicate $n = 2$ technical replicates of the $\beta$-lactamase assay performed on the same *C. testosteroni* culture at the same time.

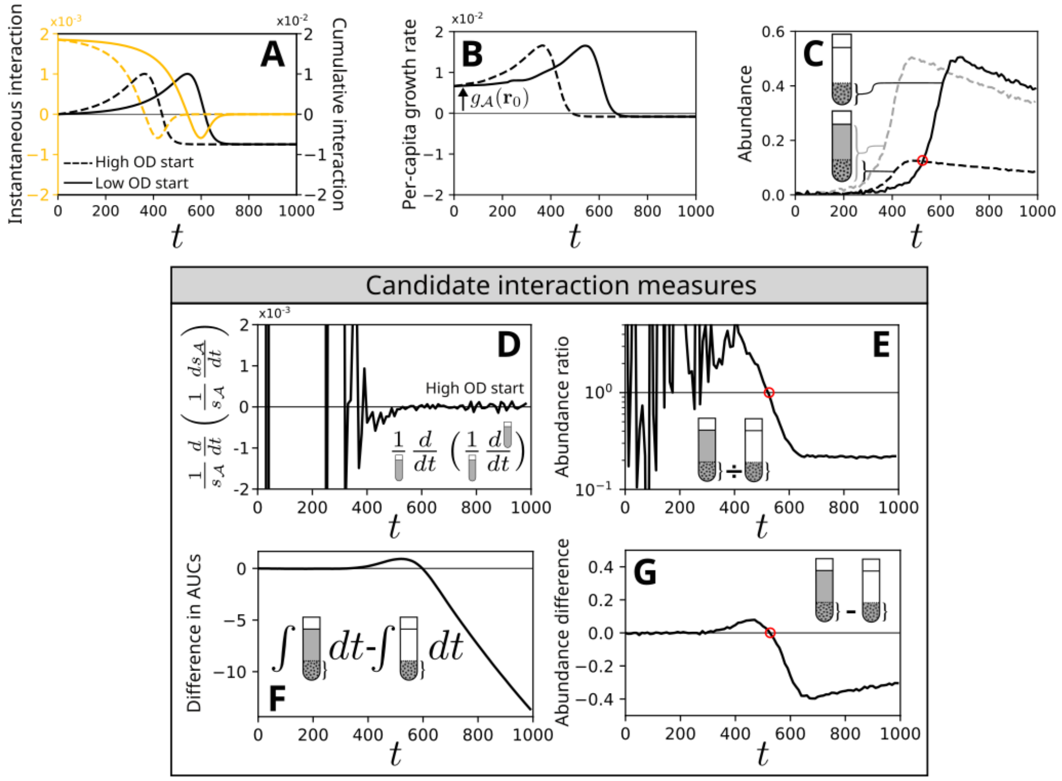


Figure S2: **Simulated batch cultures show abundance differences capture time-dependent interactions. A** We illustrate the relationship between the instantaneous, cumulative and measured intra-specific interactions for the toxin-nutrient system by simulating batch cultures initialised at high and low inoculation densities (compare Fig. 2F). **B** The per-capita growth rate of cells in each of these populations is given by adding a constant $g_\mathcal{A}(\boldsymbol{r}_0)$ – representing the growth rate of $\mathcal{A}$ in the initial environment $r_0$ – to the cumulative interaction. **C** Integration of the per-capita growth rate yields growth curves for the two conditions. Here, normally distributed noise (s.d. = 0.005) has been added to simulate measurement noise. As in Fig. 3C, we measure the size of the sub-population in the high starting-density condition (black dashed line) matched to the low starting-density population (black solid line) by dividing the high-density growth curve (gray dashed line) by the ratio of inoculation densities. **D** One option to estimate interactions from these growth curves is to calculate the quantity $\frac{1}{s_\mathcal{A}}\frac{d}{dt}\left(\frac{1}{s_\mathcal{A}}\frac{ds_\mathcal{A}}{dt}\right)$ (shown for the high inoculation-density population), which from the cEO equation is expected to return the instantaneous intra-specific interaction for monocultures. **E** Alternatively, the ratio between the sub-populations over time indicates whether the sub-population in the high starting-density culture has grown more (positive interaction) or less (negative interaction) than that in the low starting-density culture.[5,28,29] While both of these approaches work in principle, in practice measurement noise is so strongly amplified at early time points when abundances are small that useful information cannot be reliably extracted. **F** The difference between the areas under the growth curves (AUCs) from the beginning of the experiment up to a query time is another alternative.[21,30] This captures the shape of the intra-specific interaction initially, but fails to stabilise once the cumulative interaction stops changing. **G** The abundance difference[7] displays low initial noisiness and long-term stability while also capturing the overall shape of the cumulative interaction, and is the primary experimental interaction measurement we use in this manuscript.

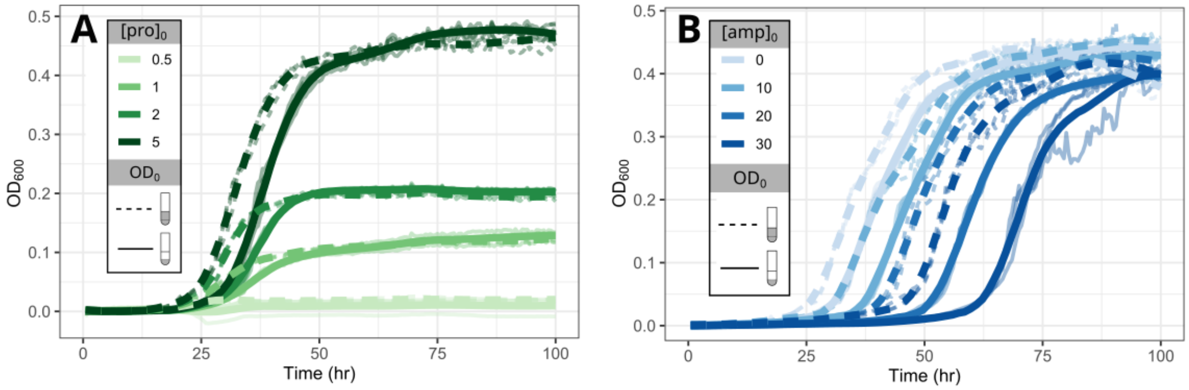


Figure S3: **Raw *C. testosteroni* growth curves under varying environmental conditions.** Raw growth curves (faint lines, $n = 3$ technical replicates for each condition) and cross-condition LOESS-smoothed averages (bold lines) for one biological replicate of the experiment shown in Fig. 3. Dashed lines indicate cultures inoculated at high OD, while solid lines indicate cultures inoculated at low OD. **A** All samples for which [amp]$_0$ $= 0$, representing the antibiotic-free control samples. **B** All samples for which [pro]$_0 = 5$mM.

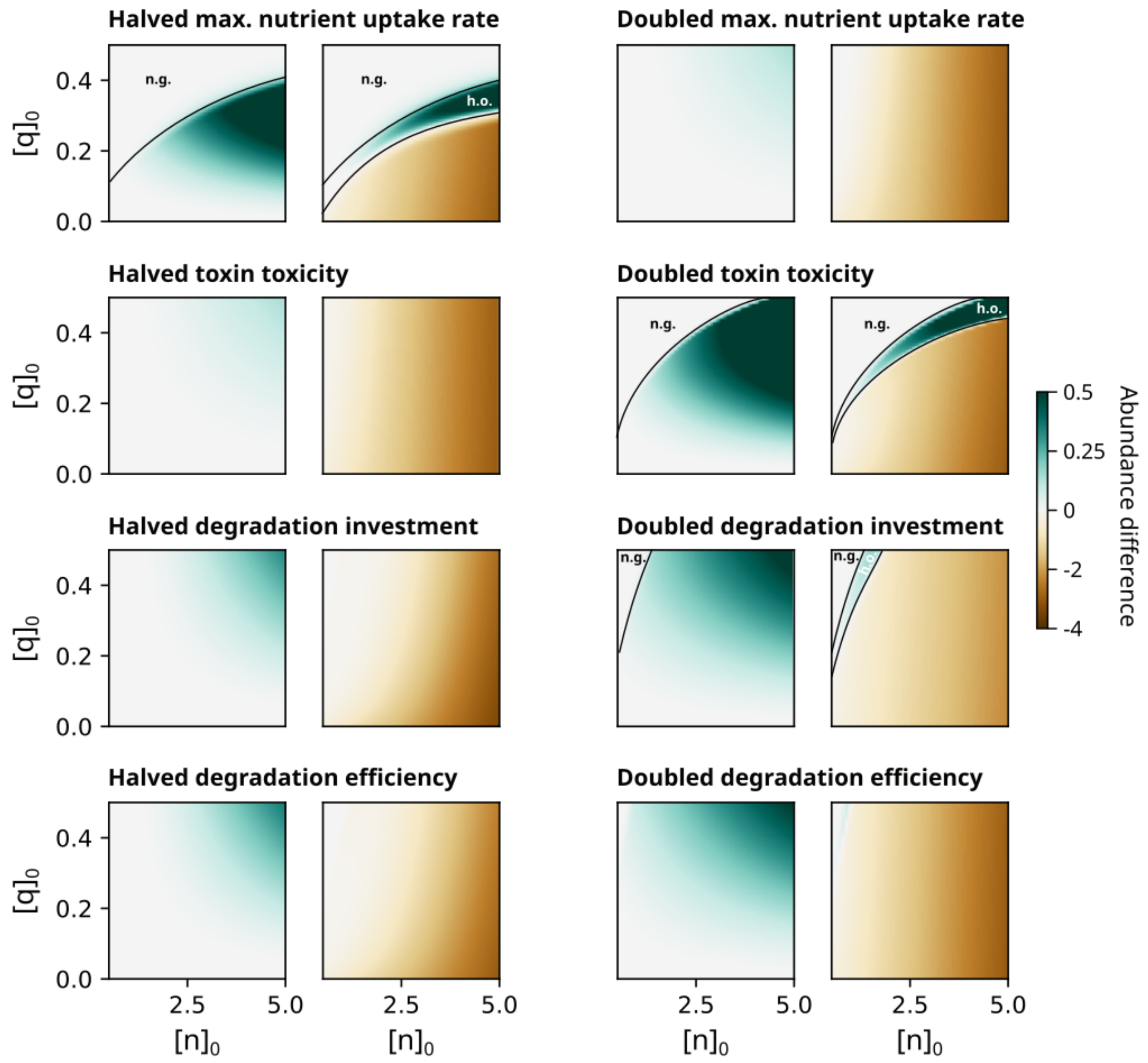


Figure S4: **Large shifts in the parameters of the toxin-nutrient model do not substantially impact the qualitative interaction patterns observed.** To investigate the robustness of the time-dependent interaction patterns observed in our toxin-nutrient model (Fig. 3H,I), we repeated our simulations with doubled and halved maximum nutrient uptake rate ($\nu_n$, top), maximal toxin impact ($\nu_q$, top-middle), toxin degradation investment ($f$, bottom-middle) and detoxification efficiency ($\delta$, bottom). In the case of doubled $\nu_q$ and halved $\nu_n$, we observe that in some environments with high toxin concentrations and low nutrient concentrations either only cells in the high inoculation OD conditions are able to grow (denoted 'h.o.') or that there is no growth for either inoculation density (denoted 'n.g.').

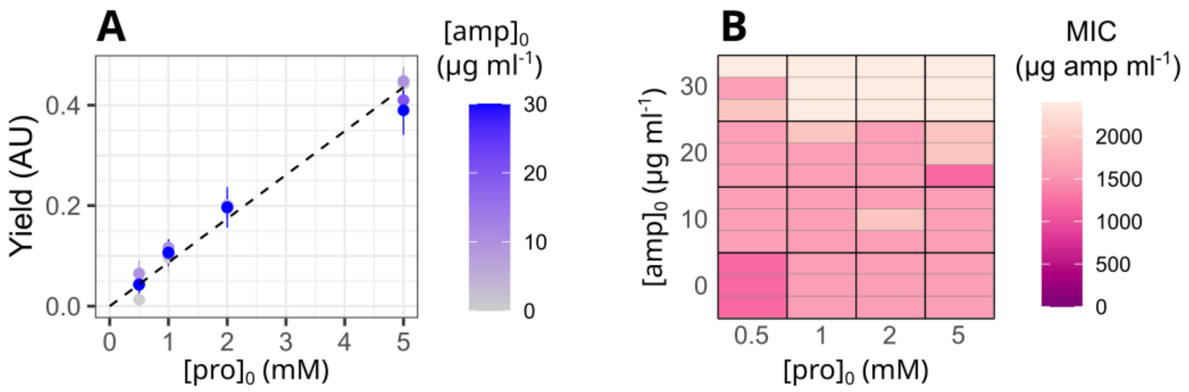


Figure S5: ***C. testosteroni* evolves stronger $\beta$-lactam resistance over experimental timescales.** Following one biological replicate of the experiment shown in Fig. 3, the minimum inhibitory concentration (MIC) of ampicillin was measured for each of the 48 low inoculation density wells (Methods). To prevent the inoculum effect from influencing our measurements,[26,27] we scaled the inoculation volume of culture by the concentration of proline in the environment. **A** This is directly proportional to the final yield of *C. testosteroni*, shown here as the OD measurement at 100 hours. This procedure thus ensured that the inoculated population size was approximately constant. **B** Heatmap showing the MIC of ampicillin measured for each environmental condition. For each condition, we show the result for each technical replicate as a separate horizontal strip.

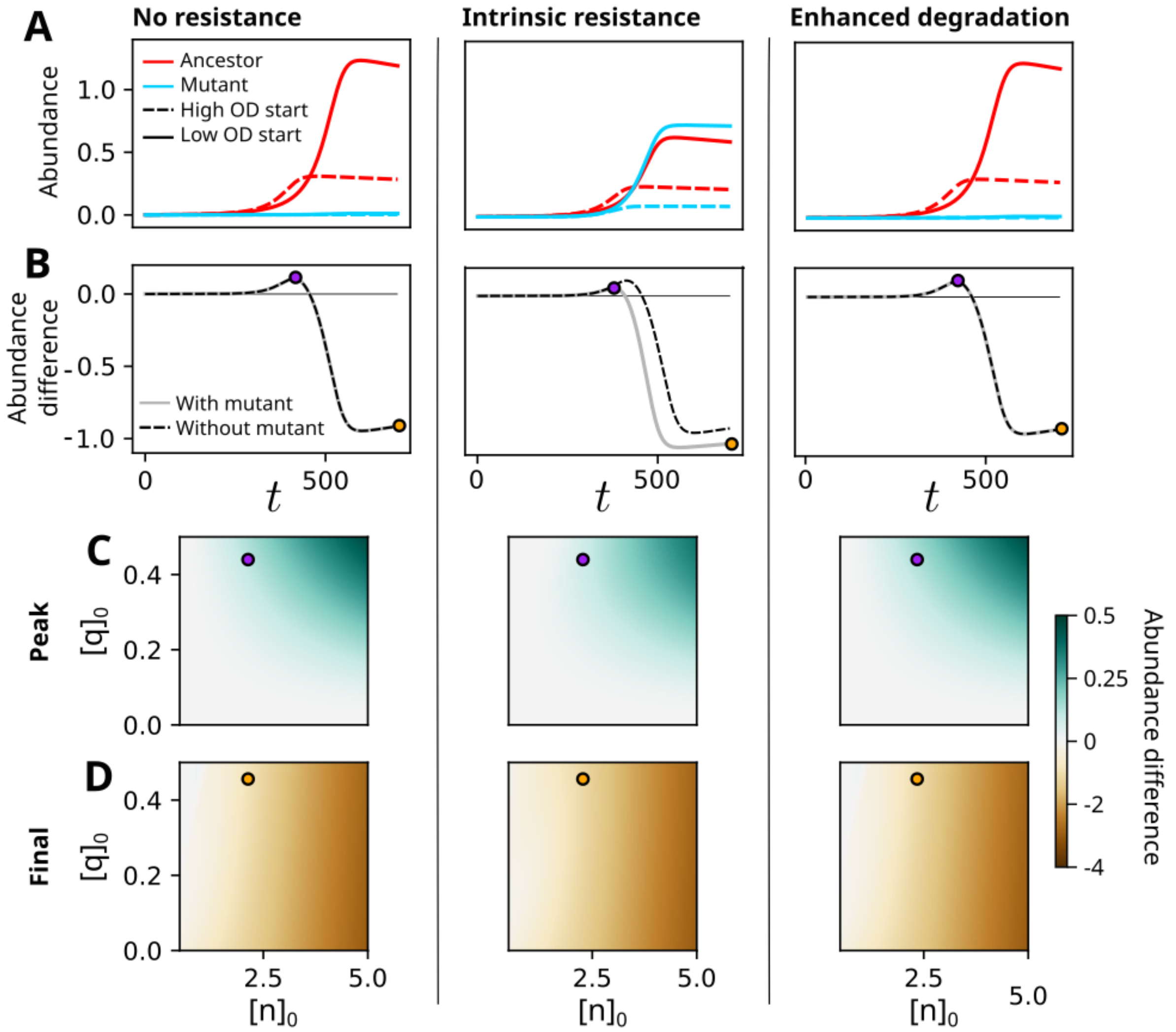


Figure S6: **Evolution of toxin resistance attenuates measured positive interaction strengths.** To investigate the effect of evolution of toxin resistance on interaction measurements in our toxin-nutrient system, we performed batch culture simulations in which an ancestral population was co-cultured with a small population (1 %) of mutant cells with either the same properties as the ancestral population (left), 5x stronger intrinsic resistance ($\nu_q/5$, middle) or 5x more efficient toxin degradation ($5\delta$, right). **A** We show raw simulated abundance curves of the ancestral (red) and mutant (blue) populations when inoculated at high (dashed lines) and low (solid lines) densities. To assist comparisons, high inoculation density curves have been normalised by the inoculation ratio (compare Fig. 3B). **B** From these, we calculate time-dependent abundance differences between the total population sizes (sum of ancestral and mutant populations) in the high and low inoculation density conditions (grey lines, compare Fig. 3C). By comparing to a null model in which only the ancestral population is present (black dashed lines), we see that only the mutant with an increase in intrinsic resistance influences the measured interaction, weakly reducing the strength of the positive phase of the measurement. **C,D** This conclusion is supported by comparing the peak (C) and final (D) abundance differences across multiple environments (compare Fig. 3H,I). Purple and orange points indicate the initial environmental composition of the simulations shown in A and B.

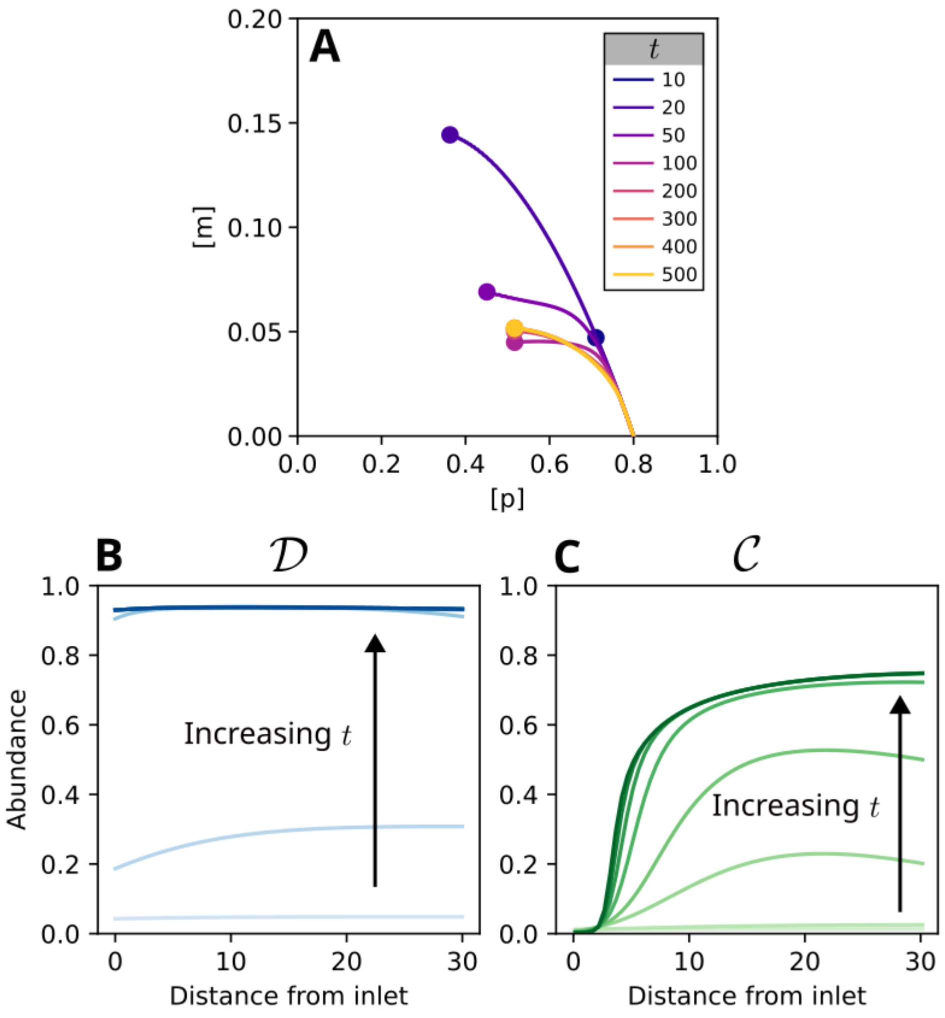


Figure S7: **Our spatial model of a microfluidic channel stabilises over long timescales.** An assumption necessary for our framework to be applied to spatially varying environments is that the composition of the environment is static (Supplementary Note 2). **A** To confirm this condition was met in our model of the microfluidic channel at long timescales, we plotted the trajectories representing the spatially-varying environment after different simulation lengths ($t$) when the flow velocity $v_x = 2.5$. These sweep out a curve from the position representing the inlet composition ($[p]_o = 0.8$, $[m]_o = 0$) to the composition at the outlet (circular points). After an initial transient during community establishment, we observe that the environmental trajectory traces out a consistent curve beyond $t = 200$. **B, C** This stabilisation of the environment corresponds to a stabilisation of the spatial structure of the community, as shown by the changing distribution of the degrader (**B**) and cross-feeder (**C**) along the length of the channel over time. Increasingly dark shades of blue and green in these plots indicate later sampling times of the simulation and correspond to the same sampling times in panel A. All other microfluidic simulations in this manuscript are sampled at $t = 1000$.

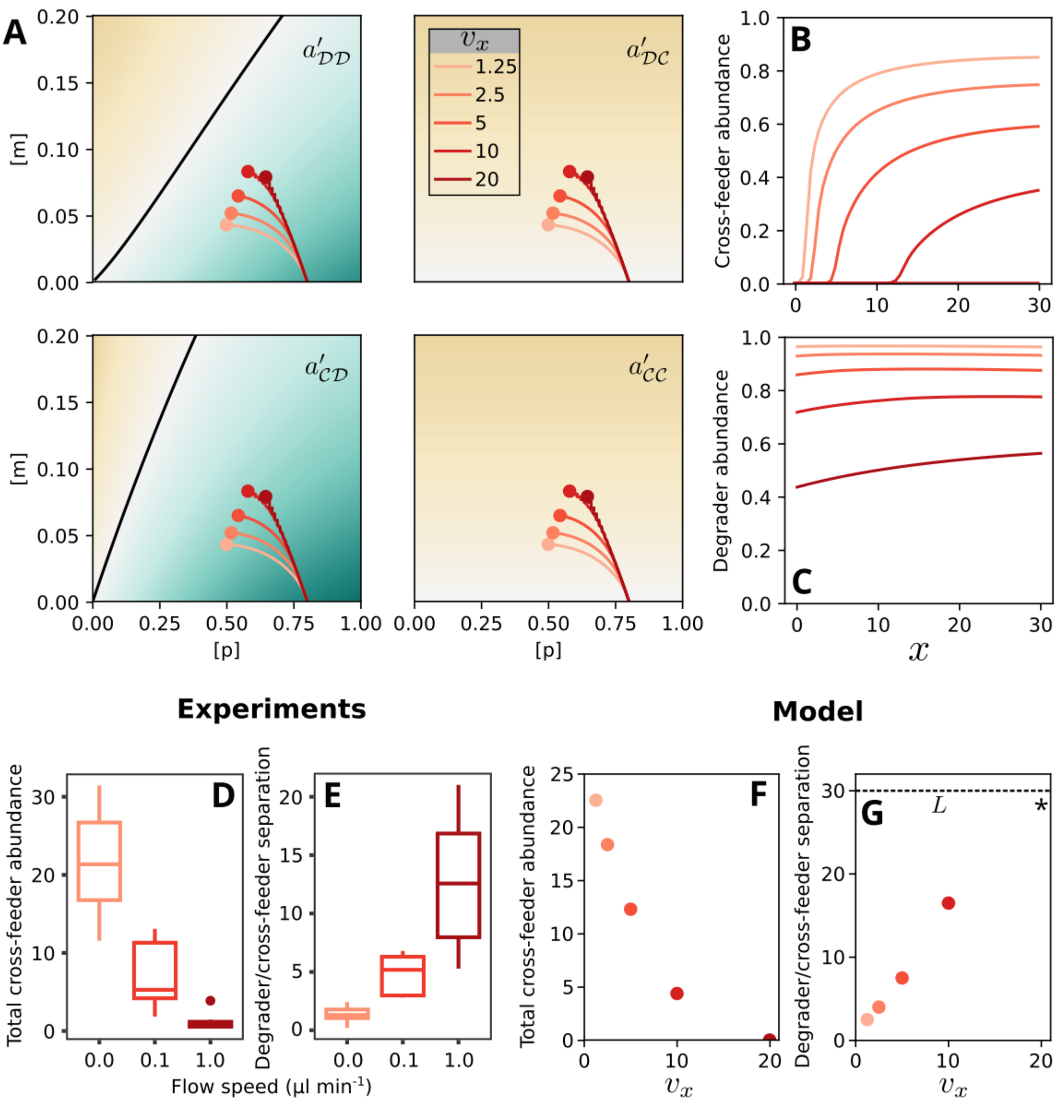


Figure S8: **Changes in the spatial structure of simulated microfluidic communities under varying flow speeds match experimental observations. A** Variation of the flow speed $v_x$ in simulated microfluidic channels results in different environmental trajectories swept out along the length of the system. Here we plot the resulting steady-state environmental trajectories on top of the instantaneous interaction maps for the degrader $\mathcal{D}$ and crossfeeder $\mathcal{C}$ species (Fig. 4D). These curves represent the variation in the composition of the environment from the inlet ($[p]_0 = 0.8$, $[m]_0 = 0$) to the outlet (circular points). **B, C** These different trajectories, combined with variations in the wash-out rate $\theta$ induced by different flow rates, result in different spatial structuring of the cross-feeder (**B**) and the degrader (**C**) along the length of the device. **D, E** The impact of flow rate on the total abundance of the cross-feeder (**D**, replotted from Wong et al.[8]) and the spatial separation between the the degrader and the cross-feeder growth zones (**E**, replotted from Wong et al.[8]) were previously quantified experimentally. **F, G** We evaluated these quantities from the steady-state species abundance profiles from our model, finding identical trends as in the experiments. The dashed line in **G** indicates the channel length $L$. As we do not observe growth of the crossfeeder in the highest flowrate simulation, we indicate that the separation between the degrader and crossfeeder growth zones is not defined for this simulation with an asterisk.

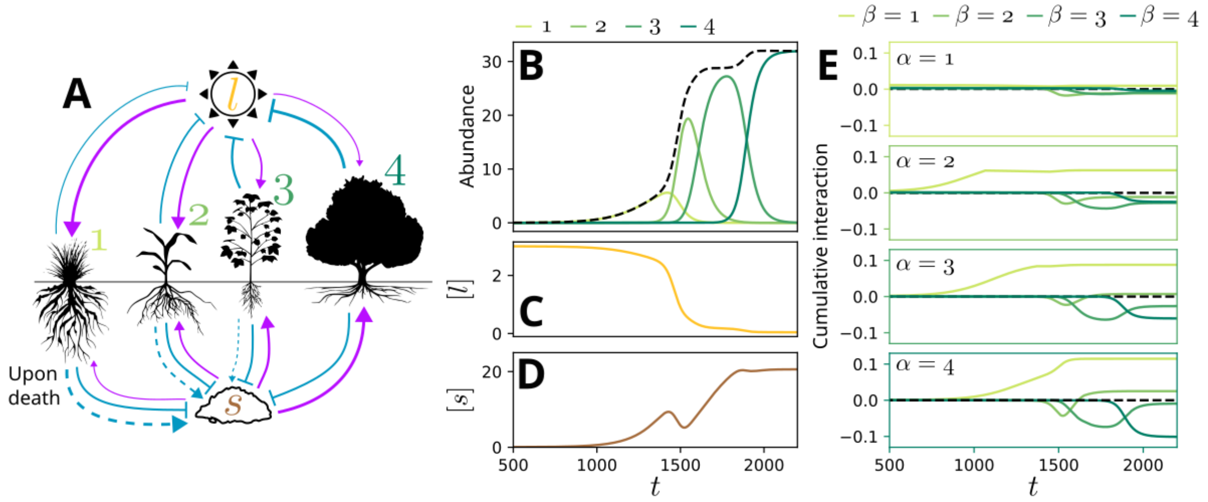


Figure S9: **Our framework can disentangle changing interactions during primary succession. A** We built an EO model of primary succession in a plant community, incorporating the resource-ratio hypothesis (Supplementary Methods). We assume a trade-off between investment into foliage and nutrient fixation, leading to a spectrum of shade-intolerant, nutrient fixing species ($1$) to shade-tolerant, soil-intensive species ($4$). Thickness of lines representing sensitivity functions (purple) indicate the necessary quantity of each resource for optimal growth, with thicker lines representing higher requirements. Thickness of lines representing uptake functions (solid blue) indicate the amount of each resource used by each species. Thickness of lines representing return functions (dashed blue) indicate the net increase in the quality of the soil when a member of the indicated species dies and decays. **B** Changes in the abundance of the four species demonstrate succession, with each of the four species giving way to the next in the successional sequence until the climax species $4$ establishes dominance. The black dashed line indicates total community abundance. **C,D** These compositional changes of the community are driven by changes in the availability of light (C) and the quality of the soil (D). **E** We can explore the changing interactions in this ecosystem by decomposing net growth rate impacts into cumulative interactions between each pair of species. Frame colour indicates the identity of the impacted species ($\alpha$), while line colour indicates the identity of the impacting species ($\beta$). Elements of panel (A) were generated with Biorender.

### 4 Supplementary tables

| Sub-component | Ingredient | Quantity |
|---|---|---|
| M9 10x | $K_2HPO_4$ | 30 g |
| | NaCl | 5 g |
| | $NH_4Cl$ | 10 g |
| | $Na_2HPO_4$ | 60 g |
| Metals 44 1000x | $Na_2EDTA \cdot 2H_2O$ | 0.387 g |
| | $ZnSO_4 \cdot 7H_2O$ | 1.095 g |
| | $FeSO_4 \cdot 7H_2O$ | 0.914 g |
| | $MnSO_4 \cdot 7H_2O$ | 0.154 g |
| | $CuSO_4 \cdot H_2O$ | 0.0392 g |
| | $Co(NO_3)_2 \cdot 6H_2O$ | 0.0284 g |
| | $Na_2B_4O_7 \cdot 10H_2O$ | 0.0177g |
| | $H_2O$ | to 1 l |
| Hutner's mineral base (HMB) 50x | Nitric triacetic acid (NTA) | 10 g |
| | $MgSO_4 \cdot 7H_2O$ | 14.45 g |
| | $CaCl_2 \cdot 2H_2O$ | 3.33 g |
| | $(NH_4)_6Mo_7O_24 \cdot 4H_2O$ | 0.00974 g |
| | $FeSO_4 \cdot 7H_2O$ | 0.099 g |
| | Metals 44 1000x | 50 ml |
| | $H_2O$ | to 1 l |
| Base minimal media | HMB 50x | 20 ml |
| | M9 10x | 100 ml |
| | $H_2O$ + proline | to 1 l |

Table 1: Composition of base minimal media.

| Symbol | Name | Value |
|---|---|---|
| $\nu_n$ | Maximal nutrient uptake rate | 0.05 |
| $\nu_q$ | Maximal toxicity | 0.05 |
| $f$ | Toxin degradation investment | 0.2 |
| $\delta$ | Toxin degradation efficiency | 10 |

Table 2: Parameter values used for the toxin-nutrient model.

| Symbol | Name | Value |
|---|---|---|
| $\nu_{\mathcal{D}p}$ | Maximal polymer consumption rate | 1 |
| $\nu_{\mathcal{D}m}$ | Maximal monomer uptake rate (degrader) | 1 |
| $\nu_{\mathcal{C}m}$ | Maximal monomer uptake rate (cross-feeder) | 1 |
| $Y_{\mathcal{D}m}$ | Monomer yield (degrader) | 1 |
| $Y_{\mathcal{C}m}$ | Monomer yield (cross-feeder) | 1 |
| $K_{\mathcal{D}m}$ | Half-velocity constant of monomer uptake (degrader) | 1 |
| $K_{\mathcal{C}m}$ | Half-velocity constant of monomer uptake (cross-feeder) | 1 |
| $\phi$ | Polymer to monomer conversion fraction | 0.6 |

Table 3: Parameter values used for the degrader-crossfeeder model.

| Symbol | Name | Values ($\mathcal{X} \in \{1, 2, 3, 4\}$) |
|---|---|---|
| $\nu_{\mathcal{X}}$ | Maximal population growth rate | 0.1, 0.1, 0.1, 0.1 |
| $\theta_{\mathcal{X}}$ | Mortality rate | 0.03, 0.03, 0.03, 0.03 |
| $\zeta_{\mathcal{X}}$ | Fractional light reduction by one unit of $\mathcal{X}$ | 0.1, 0.2, 0.3, 0.4 |
| $Y_{\mathcal{X}g}$ | Size of population $\mathcal{X}$ that depletes one unit of soil quality | 1, 1, 1, 1 |
| $Y_{\mathcal{X}d}$ | Soil enrichment from decomposition of one unit of $\mathcal{X}$ | 1.45, 1.15, 1.05, 1 |
| $K_{\mathcal{X}s}$ | Half-velocity constant of soil use | 0.1, 0.4, 1.25, 5 |
| $K_{\mathcal{X}l}$ | Half-velocity constant of light use | 5, 1.25, 0.4, 0.1 |
| $L_{\text{max}}$ | Maximal light availability | 3 |

Table 4: Parameter values used for the successional model.

### References


[1] Novak, M.; Yeakel, J. D.; Noble, A. E.; Doak, D. F.; Emmerson, M.; Estes, J. A.; Jacob, U.; Tinker, M. T.; Wootton, J. T. *Annual Review of Ecology, Evolution, and Systematics* **2016**, *47*, 409–432.

[2] Meszéna, G.; Gyllenberg, M.; Pásztor, L.; Metz, J. A. *Theoretical Population Biology* **2006**, *69*, 68–87.

[3] Koffel, T.; Daufresne, T.; Klausmeier, C. A. *Ecological Monographs* **2021**, *91*, e01458.

[4] Mitri, S.; Richard Foster, K. *Annual Review of Genetics* **2013**, *47*, 247–273, PMID: 24016192.

[5] Kehe, J.; Ortiz, A.; Kulesa, A.; Gore, J.; Blainey, P. C.; Friedman, J. *Science Advances* **2021**, *7*, 7159.

[6] Daniels, M.; van Vliet, S.; Ackermann, M. *The ISME Journal 2023* **2023**, *17*, 1–11.

[7] Foster, K. R.; Bell, T. *Current Biology* **2012**, *22*, 1845–1850.

[8] Wong, J. P. H.; Fischer-Stettler, M.; Zeeman, S. C.; Battin, T. J.; Persat, A. *Proceedings of the National Academy of Sciences of the United States of America* **2023**, *120*, e2217577120.

[9] Roberts, D. W. *Vegetatio* **1987**, *69*, 27–33.

[10] Butterfield, B. J.; Callaway, R. M. *Functional Ecology* **2013**, *27*, 907–917.

[11] Tansley, A. G. *Ecology* **1935**, *16*, 284–307.

[12] Connell, J. H.; Slatyer, R. O. *The American Naturalist* **1977**, *111*, 1119–1144.

[13] McCook, L. J. *Vegetatio* **1994**, *110*, 115–147.

[14] del Moral, R.; Thomason, L. A.; Wenke, A. C.; Lozanoff, N.; Abata, M. D. *Journal of Vegetation Science* **2012**, *23*, 73–85.

[15] Walker, L. R.; Clarkson, B. D.; Silvester, W. B.; Clarkson, R., Beverley *Journal of Vegetation Science* **2003**, *14*, 277–290.

[16] Pacala, S.; Rees, M. *The American Naturalist* **1998**, *152*, 729–737.

[17] Tilman, D. *The American Naturalist* **1985**, *125*, 827–852.

[18] Chapin, F. S.; Walker, L. R.; Fastie, C. L.; Sharman, L. C. *Ecological Monographs* **1994**, *64*, 149–175.

[19] Morris, W. F.; Wood, D. M. *Ecology* **1989**, *70*, 697–703.

[20] Berkowitz, A. R.; Canham, C. D.; Kelly, V. R. *Ecology* **1995**, *76*, 1156–1168.

[21] Piccardi, P.; Vessman, B.; Mitri, S. *Proceedings of the National Academy of Sciences of the United States of America* **2019**, *116*, 15979–15984.

[22] Drescher, K.; Nadell, C. D.; Stone, H. A.; Wingreen, N. S.; Bassler, B. L. *Current Biology* **2014**, *24*, 50–55.

[23] Sulheim, S.; Mitri, S. *Trends in Microbiology* **2023**, *31*, 426–427.

[24] Smith, H. L.; Waltman, P. *The Theory of the Chemostat: Dynamics of Microbial Competition*; Cambridge University Press, 1995.

[25] Kim, J.; Kim, H.-S.; Han, S.; Lee, J.-Y.; Oh, J.-E.; Chung, S.; Park, H.-D. *Lab on a Chip* **2013**, *13*, 1846–1849.

[26] Parker, R. F. *Experimental Biology and Medicine* **1946**, *63*, 443–446.

[27] Lenhard, J. R.; Bulman, Z. P. *Journal of Antimicrobial Chemotherapy* **2019**, *74*, 2825–2843.

[28] Hsu, R. H.; Clark, R. L.; Tan, J. W.; Ahn, J. C.; Gupta, S.; Romero, P. A.; Venturelli, O. S. *Cell Systems* **2019**, *9*, 229–242.e4.

[29] Vos, M. G. D.; Zagorski, M.; McNally, A.; Bollenbach, T. *Proceedings of the National Academy of Sciences of the United States of America* **2017**, *114*, 10666–10671.

[30] Weiss, A. S. et al. *The ISME Journal 2021 16:4* **2021**, *16*, 1095–1109.
